# Humatch - fast, gene-specific joint humanisation of antibody heavy and light chains

**DOI:** 10.1101/2024.09.16.613210

**Authors:** Lewis Chinery, Jeliazko R. Jeliazkov, Charlotte M. Deane

**Affiliations:** University of Oxford, Research Technologies, GSK R&D; Protein Design and Informatics, Research Technologies, GSK R&D

## Abstract

1

Antibodies are a popular and powerful class of therapeutic due to their ability to exhibit high affinity and specificity to target proteins. However, the majority of antibody therapeutics are not genetically human, with initial therapeutic designs typically obtained from animal models. Humanisation of these precursors is essential to reduce immunogenic risks when administered to humans.

Here, we present Humatch, a computational tool designed to offer experimental-like joint humanisation of heavy and light chains in seconds. Humatch consists of three lightweight Convolutional Neural Networks (CNNs) trained to identify human heavy V-genes, light V-genes, and well-paired antibody sequences with near-perfect accuracy. We show that these CNNs, alongside germline similarity, can be used for fast humanisation that aligns well with known experimental data. Throughout the humanisation process, a sequence is guided towards a specific target gene and away from others via multiclass CNN outputs and gene-specific germline data. This guidance ensures final humanised designs do not sit ‘between’ genes, a trait that is not naturally observed. Humatch’s optimisation towards specific genes and good VH/VL pairing increases the chances that final designs will be stable and express well and reduces the chances of immunogenic epitopes forming between the two chains. Humatch’s training data and source code are provided open-source.

**Availability:** Source code is freely available at github.com/oxpig/Humatch. Data can be found at doi.org/10.5281/zenodo.13764770

**Supplementary information:** Supplementary data are available at *bioRxiv* online.

## 2 Introduction

The antibody drug discovery process is a challenging, multi-objective optimisation problem. This problem requires the development of potential leads that bind their target strongly (high affinity), have few off-target effects (high specificity), and possess good developability characteristics[1].

One critical step in the development of antibody therapeutics is humanisation[2]. Humanisation is important as in many instances, drug precursors originate in animal models. Gordon *et al.*[3] found that approximately 60% of therapeutics listed in Thera-SAbDab[4] are not genetically human in origin and that this percentage has remained constant over the past two decades.

In humanisation workflows, animals, such as mice, are exposed to the antigen of interest, an immune response is raised, and dominant clones are obtained through library screens[5]. These clones, which constitute precursor therapeutics, have binding sites optimised to bind the target antigen. However, the rest of the antibody, predominantly the framework region (FR), could contain human immunogenic epitopes. These epitopes risk raising anti-drug antibody (ADA) responses in human patients[2]. It is therefore critical to mutate these regions before starting human trials whilst maintaining strong binding and high expression.

Classical humanisation techniques may involve grafting antigen-specific Complementarity Determining Region loops (CDRs) onto a human antibody framework and back-mutating Vernier zone residues to the precursor sequence[6, 7]. Alternatively, iterative mutations towards a target human germline are made, typically on surface-accessible residues, with these optimised through experimental trial and error[8, 9]. These classical approaches can succeed in humanising precursor sequences but are time and cost-intensive, require many mutations that could disrupt binding, and may still lead to therapeutics with high ADA levels[10]. Recently, computational tools have been developed to aid in this process.

Hu-mAb[10] is one such computational tool that consists of many human gene-specific random forest (RF) classifiers. These RFs were trained on data from the Observed Antibody Space (OAS)[11, 12] database and succeeded in identifying human heavy and light V-genes with near 100% accuracy on Hu-mAb’s test set. Hu-mAb humanises heavy and light chains separately by making all possible single-point mutations to a starting sequence, scoring them with its RF models, selecting the top variant, and repeating until a target threshold is met. Hu-mAb is widely used, however, the humanisation process is slow (~18 minutes) and susceptible to getting stuck in local minima. Furthermore, its classifiers were only trained on species where OAS contained significant sequence data and some human V-genes are missing. CDR mutations are also strictly forbidden.

BioPhi[13] is an alternative platform consisting of OASis, a humanness classifier and Sapiens, a transformer-based[14] humanisation tool. OASis compares all possible 9-mers in an input sequence to how often each is observed within a set of human sequences from OAS. 9-mers that are observed frequently are ranked highly, while those rarely observed receive low scores. Sapiens humanises sequences towards high OASis scores and is trained solely on OAS human sequences using a masked language model approach. During humanisation, probabilities for each residue position are calculated and the amino acids with top predictions are selected (CDR residues are ignored). This process is repeated up to four times. Given all probabilities are calculated in one single pass, humanisation is fast. However, Sapiens lacks the option to select the desired germline; instead, this selection is implicit during training. Additionally, OASis’ 9-mer peptide scoring system can rank sequences that are ‘between’ genes highly e.g. starting HV1 and ending HV3. Finally, like Hu-mAb, Sapiens optimises heavy and light chains independently. This separate humanisation risks disrupting expression, lowering stability, and creating immunogenic epitopes that span both chains[15, 16].

AbNatiV[17], like Sapiens, was trained with masked unsupervised learning on human antibody sequences from OAS. Three antibody (and one nanobody) models were trained, each a vector-quantised variational auto-encoder, for heavy (HV), kappa (KV), and lambda (LV) sequences. Nativeness scores (masked likelihoods) were then calculated per residue and averaged to provide an overall nativeness score. This nativeness score can be used to classify human from non-human sequences with high accuracy. The humanisation process is similar to Sapiens, though only mutations to residues observed in its human training data (calculated using a position-specific scoring matrix, PSSM) are allowed by default. Mutations are also limited to solvent-accessible framework residues. This step seeks to avoid unnecessary mutations to residues that cannot form immunogenic epitopes, however, the structural modelling required increases run time. To lessen this time, a single model of the wildtype structure is used for all variants but this model risks becoming unrepresentative as the number of mutations increases. Also, the PSSM used during humanisation is not germline-specific, potentially allowing mutations to hybrid mixed genes, similar to Sapiens. AbNatiV, like Hu-mAb and Sapiens, humanises the chains independently.

In this paper, we describe a fast and accurate human classification and humanisation tool, Humatch. The humanisation logic of Humatch produces sequences that align well with experimentally optimised sequences. Humatch is also the first humanisation tool to jointly humanise heavy and light chains with the goal of maintaining high expression, good stability, and removing potential immunogenic epitopes formed across the two chains. Furthermore, Humatch is designed to consistently push designs towards single human V-genes at all stages, avoiding potential hybrid mixed gene designs. Humatch is available open-source and can be easily integrated into existing development pipelines.

## 3 Results

### 3.1 Humatch classifies human heavy, light, and naturally paired sequences with high accuracy

Humatch is a set of three Convolutional Neural Networks (CNNs). The first two are trained on unpaired human and non-human antibody sequences; one on heavy sequences only (CNN-H), and one on light sequences (CNN-L). Sequences are padded to the most common 200 IMGT[18] positions identified by KASearch[19] and amino acids are represented by 10-dimensional Kidera[20] feature vectors. Full details on data preparation and model architecture can be found in the Methods section. Both CNNs output a multiclass prediction vector that contains the probability the input sequence is either non-human or one of the human V-genes (HV1-7 for heavy sequences or LV1-10 and KV1-7 for light sequences). The class predictions for each classifier are softmaxed and sum to one.

When trained on aligned sequence data from the Observed Antibody Space (OAS)[11, 12] database (see Methods), the two CNNs are able to separate all human V-genes from one another and non-human sequences with high accuracy (Table 1). Some V-genes, such as KV7, have lower accuracy than other classes (PR AUC of 0.85 vs 0.99+ for all other classes) due to a lack of training data (4,000 sequences vs 3m+ for the most populous class, see Methods). In our Supplementary Information (SI) we show Humatch maintains high accuracy even when the training and test data is split by allele.

**Table 1:**
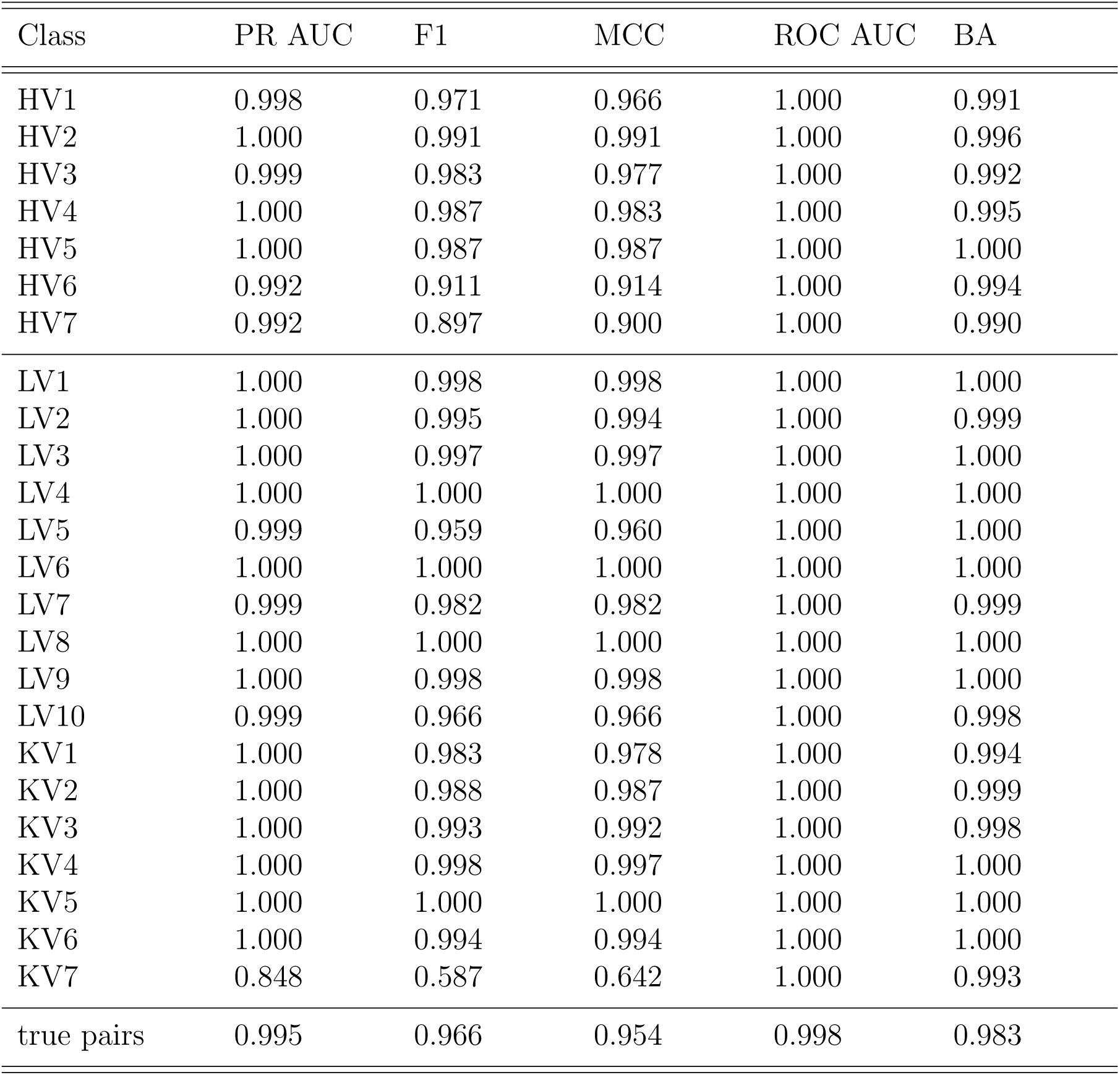
Performance of Humatch’s three CNN classifiers - heavy, light, and paired. Sequences belonging to all genes are classified with high accuracy. Gene classes with the lowest scores tend to have little training data available. PR AUC = Area Under the Precision-Recall Curve; F1 = F1-score, MCC = Matthews Correlation Coefficient; ROC AUC = Area Under the Receiver Operating Characteristic Curve; BA = Balanced Accuracy.

| Class | PR AUC | F1 | MCC | ROC AUC | BA |
| --- | --- | --- | --- | --- | --- |
| HV1 | 0.998 | 0.971 | 0.966 | 1.000 | 0.991 |
| HV2 | 1.000 | 0.991 | 0.991 | 1.000 | 0.996 |
| HV3 | 0.999 | 0.983 | 0.977 | 1.000 | 0.992 |
| HV4 | 1.000 | 0.987 | 0.983 | 1.000 | 0.995 |
| HV5 | 1.000 | 0.987 | 0.987 | 1.000 | 1.000 |
| HV6 | 0.992 | 0.911 | 0.914 | 1.000 | 0.994 |
| HV7 | 0.992 | 0.897 | 0.900 | 1.000 | 0.990 |
| LV1 | 1.000 | 0.998 | 0.998 | 1.000 | 1.000 |
| LV2 | 1.000 | 0.995 | 0.994 | 1.000 | 0.999 |
| LV3 | 1.000 | 0.997 | 0.997 | 1.000 | 1.000 |
| LV4 | 1.000 | 1.000 | 1.000 | 1.000 | 1.000 |
| LV5 | 0.999 | 0.959 | 0.960 | 1.000 | 1.000 |
| LV6 | 1.000 | 1.000 | 1.000 | 1.000 | 1.000 |
| LV7 | 0.999 | 0.982 | 0.982 | 1.000 | 0.999 |
| LV8 | 1.000 | 1.000 | 1.000 | 1.000 | 1.000 |
| LV9 | 1.000 | 0.998 | 0.998 | 1.000 | 1.000 |
| LV10 | 0.999 | 0.966 | 0.966 | 1.000 | 0.998 |
| KV1 | 1.000 | 0.983 | 0.978 | 1.000 | 0.994 |
| KV2 | 1.000 | 0.988 | 0.987 | 1.000 | 0.999 |
| KV3 | 1.000 | 0.993 | 0.992 | 1.000 | 0.998 |
| KV4 | 1.000 | 0.998 | 0.997 | 1.000 | 1.000 |
| KV5 | 1.000 | 1.000 | 1.000 | 1.000 | 1.000 |
| KV6 | 1.000 | 0.994 | 0.994 | 1.000 | 1.000 |
| KV7 | 0.848 | 0.587 | 0.642 | 1.000 | 0.993 |
| true pairs | 0.995 | 0.966 | 0.954 | 0.998 | 0.983 |

Other humanness classifiers, such as Hu-mAb[10], report higher accuracy metrics. However, Hu-mAb’s training data did not include harder-to-separate rhesus sequences, or the data sparse V-genes LV9 and KV7. Humatch can achieve comparable accuracy to Hu-mAb when trained and tested on the same data (see SI). To prevent overfitting and allow for a smoother humanisation process, we stopped our CNN training early after three epochs for both CNN-H and CNN-L.

In addition to unpaired heavy and light chain classifiers, Humatch includes a third CNN (CNN-P) trained on naturally paired human heavy and light chains and artificially ‘badly’ paired sequences. Training of CNN-P was also stopped early after 15 epochs to prevent overfitting. The artificially ‘badly’ paired sequences used in training the third CNN were created by joining heavy and light chains from different B-cell classes, such as memory and naive, and from different studies in OAS (see Methods). This criteria for artificial pairing means that the chains are unlikely, but not guaranteed, to match poorly. Despite potential noise in the training data that arises from the use of artificial bad pairing, Humatch can accurately separate naturally paired from artificially ‘badly’ paired human sequences (PR AUC and ROC AUC above 0.99, see Table 1).

CNN-P is designed to ensure that the variable heavy (VH) and variable light (VL) regions remain well-matched during the humanisation process. This pairing consideration aims to maintain good expression and stability and account for the fact that immunogenic epitopes can be formed across the two chains[15, 16]. To demonstrate the utility of including a paired model we examined the relationship between high CNN-P predictions and thermal stability using 137 therapeutics with melting temperature data from Jain *et al.*[21] (see SI). On average, we found that those with high CNN-P predictions (above 0.5) had higher melting temperatures than those with lower predictions (*p <* 0.001).

Humatch as standard uses the outputs of all three CNNs for paired VH/VL humanness classification, though the unpaired models can be run individually. Default thresholds of 0.95 are recommended for all classifiers to ensure high precision and recall (see SI). These thresholds can all be adjusted by users.

### 3.2 Humatch identifies therapeutics that are more likely to have high anti-drug antibody responses

Marks *et al.*[10] introduced a dataset of 217 therapeutics where the anti-drug antibody (ADA) levels had been recorded and corresponding sequence data was found for 211 of them. These sequences were numbered and aligned using ANARCI[22] and Humatch was then used to obtain heavy, light, and paired predictions for all 211 therapeutics.

The minimum score of the three CNNs was calculated, reasoning that if one of the CNNs ranked a therapeutic poorly, then it should be classed as higher risk. The minimum of the CNN predictions was found to correlate with ADA response (Figure 1), with the majority of highly immunogenic therapeutics having low Humatch predictions. Using a loose Humatch cut-off of 0.01 to retain most low ADA therapeutics removed 89% of high ADA (red), 59% of medium ADA (yellow), and just 23% of low ADA (green) therapeutics. Alternative humanness classifiers, including Hu-mAb, AbNatiV, and OASis, offer similar separation of low and high ADA antibodies (see SI). Further breakdowns of ADA responses for Humatch’s three classifiers can also be found in the SI.

**Figure 1:**
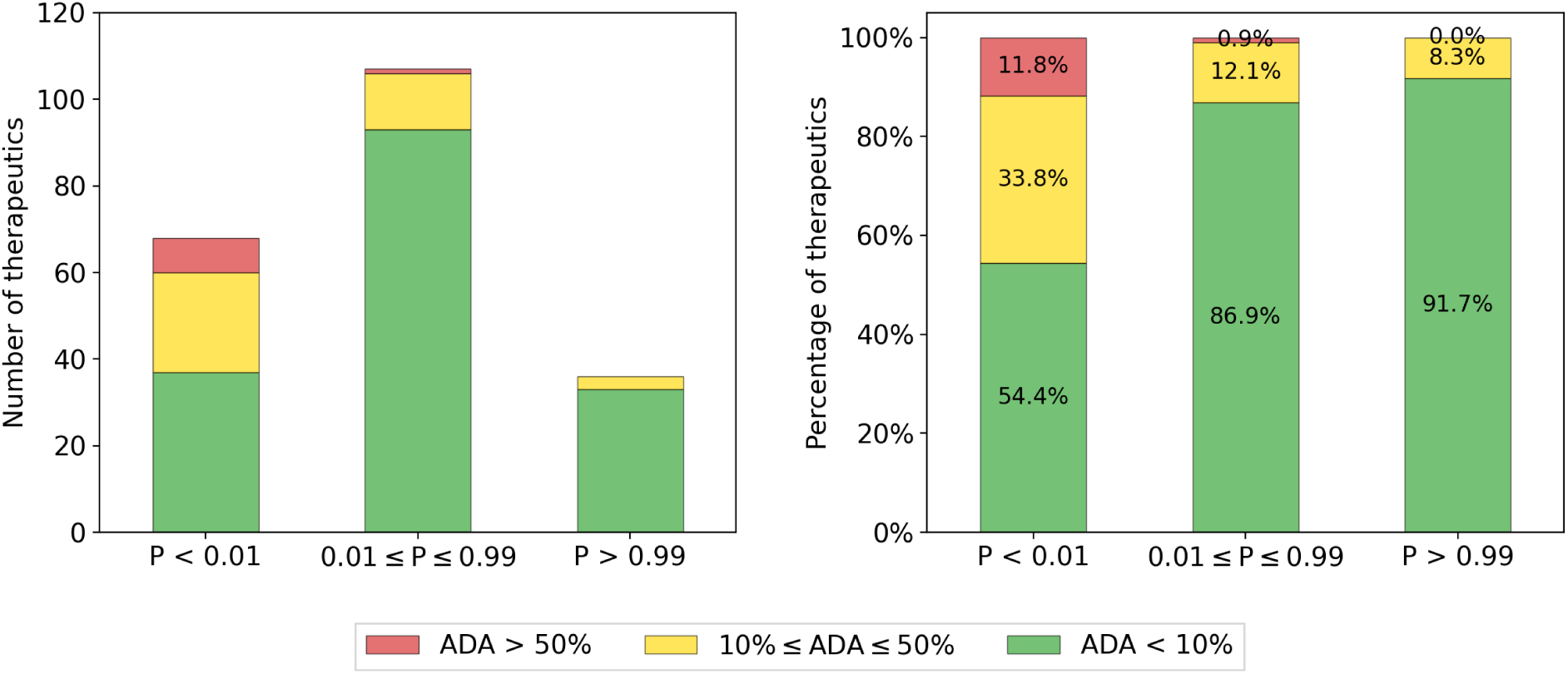
Categorisation of 211 therapeutics with anti-drug antibody labels by Humatch. The 211 therapeutics were scored by Humatch’s heavy, light, and paired CNNs. The minimum of these three scores (*P*) was used to group the therapeutics into three bins - those with a minimum prediction above 0.99, below 0.01, or between these values. The left plot shows the total number of therapeutics that were grouped in each bin and the right plot shows the proportions broken down by bin. The majority of therapeutics with the highest ADA levels (*>*50%) fall in the lowest prediction bin (*P <* 0.01).

### 3.3 Humatch-humanised sequences align well with experiments

Humatch is primarily designed to offer experimental-like humanisation in seconds. This humanisation is achieved through a combination of rapid gene-specific germline-lookup tables and guidance by Humatch’s three CNNs. Figure 2 shows an overview of all the main steps involved in the humanisation process and these are summarised below. Further details are provided in the Methods.

**Figure 2:**
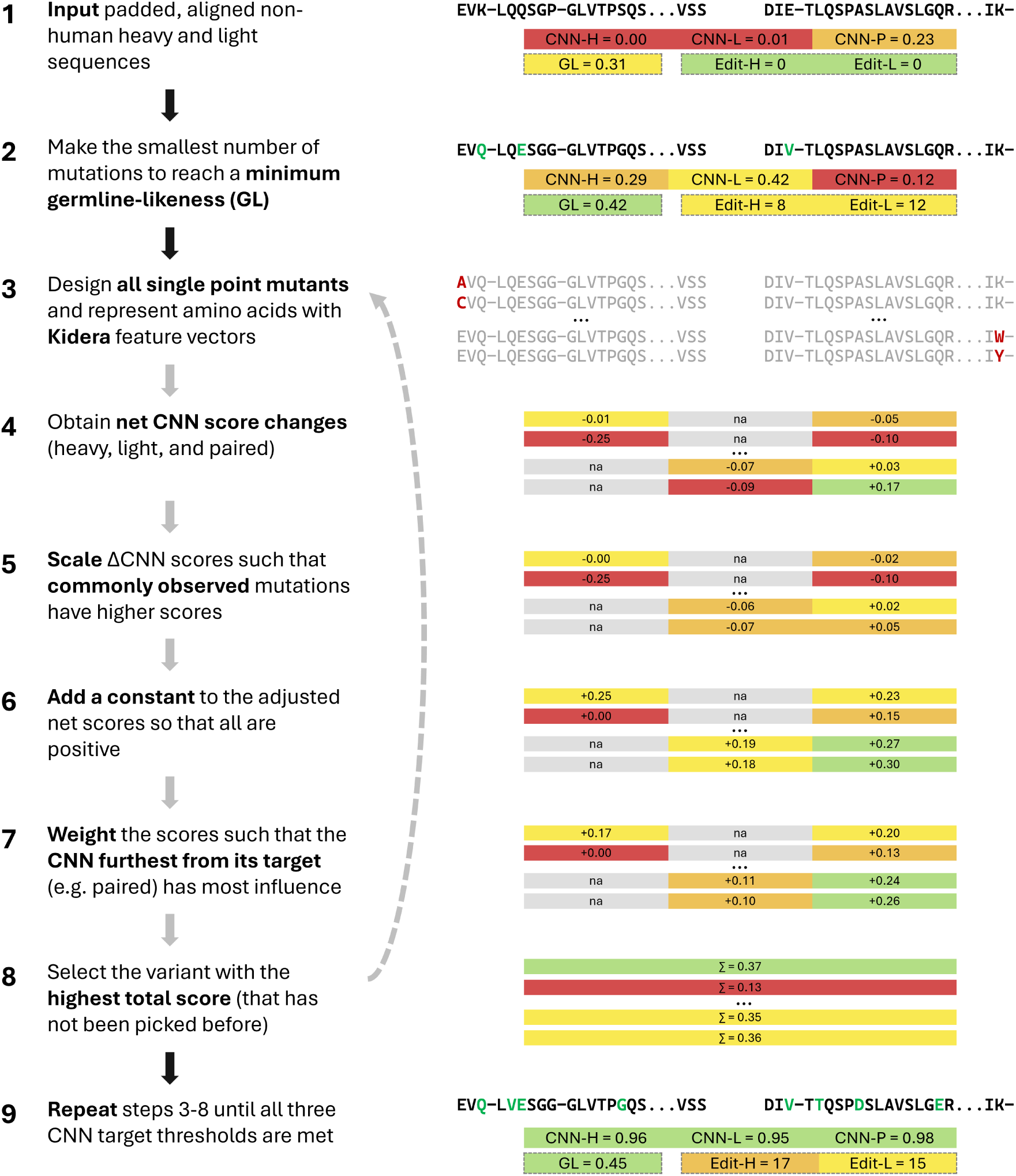
An overview of Humatch’s humanisation procedure including example sequences, CNN scores (H=heavy, L=light, P=paired), germline-likeness scores (GL), and heavy and light precursor edit distances. All numbers are for demonstration purposes only. If at any point during the humanisation process all CNN target thresholds (default 0.95) are met the humanisation process exits. Targeting an initial germline-likeness (step 2) is fast and places Humatch on a sensible humanisation trajectory. All possible single-point variants are designed in step 3, but only four are included in the figure for clarity. Users may select residues to remain unmutated throughout the process (CDRs are ignored by default). In step 8, if the highest-scoring sequence has been selected before, the next highest-scoring novel sequence is selected, avoiding Humatch becoming stuck in local minima. The humanisation process can exit early if a total maximum allowed edit distance is reached (default 60).

The first step in the humanisation process involves the most common germline mutations being made to the input sequence to achieve a minimum germline-likeness (GL) score of 0.40 for both heavy and light chains (see Methods and SI). This initial GL-matching step is fast and allows the humanisation of a starting sequence to any target gene. Next, all single-point variants of the GL-mutated sequence are designed and scored with Humatch’s three CNNs. These scores are then scaled to up-weight common germline mutations and prioritise improvements to the lowest CNN score (see Methods and SI). The top-scoring variant at this stage is selected and this process is repeated until all three CNN target thresholds of 0.95 are met. In successive iterations, Humatch ignores previous top-scoring variants to avoid local minima. Target GL scores of 0.40 and CNN thresholds of 0.95 were decided based on observed therapeutics GL scores and desired CNN precision-recall values (see SI).

To test Humatch we used it to humanise a set of 25 precursor heavy and light sequences with known experimentally humanised therapeutic endpoints[10]. The target scores were set for Humatch following a similar process to Hu-mAb, using the higher of 0.95 or the scores achieved by the experimentally optimised sequences (multiplied by 0.9999 to avoid forcing CNN scores to 1). All sequences reached their humanisation target scores in fewer than 60 edits (the maximum observed experimentally).

We also conducted a baseline experiment where only the top germline mutations were made to test the importance of Humatch’s CNN guidance in the humanisation process. In this baseline, the same number of heavy and light chain mutations were made as in Humatch’s full humanisation logic. We found that no baseline-designed paired sequences matched Humatch’s designs exactly and only two designs passed all three CNN target thresholds of 0.95 (further details can be found in the SI).

Table 2 compares Humatch’s humanisation performance to three other antibody humanisation tools - Hu-mAb[10], Sapiens[13], and AbNatiV[17]. Hu-mAb’s overlap values and edit distances were taken from their paper. All other tools were run with default and paper-recommended parameters (see SI). We observed high overlap in the mutations made by Humatch and those made experimentally. Table 2 shows that, on average, 77% of heavy and 82% of light chain mutations exactly matched between the two, higher than all other methods. The majority of these mutations were to common germline residues. Successful humanisation, according to Humatch’s three CNNs, can be achieved in fewer edits by setting a lower initial target germline-likeness score at the expense of slightly smaller overlaps and longer humanisation runtimes (see SI). These parameters can be changed by the user when running Humatch as desired.

**Table 2:** A summary of the outputs of four different humanisation tools when tasked with humanising the same 25 precursor therapeutics. Computationally designed sequences were deemed humanised when their respective target scores were met. These designs are compared to the experimentally optimised sequences (bottom). ‘H edit’ and ‘L edit’ are the mean edit distances of the humanised therapeutics from their corresponding precursor sequences. ‘H overlap’ and ‘L overlap’ described the mean overlap in mutations made computationally compared to those made experimentally. If all suggested computational mutations were made experimentally, the overlap would be one; if none matched, the overlap would be zero. ‘Time (s)’ is the mean time in seconds required to humanise each therapeutic (both heavy and light chains). The mean time required for experimental humanisation is not known. The ‘**best**’ computational scores are in bold and *second best* in italics.

| Method | H overlap | L overlap | H edit | L edit | Time (s) |
| --- | --- | --- | --- | --- | --- |
| Hu-mAb | *0.67* | *0.77* | *14.8* | *10.6* | 1,093 |
| AbNatiV | 0.57 | 0.65 | **11.5** | **9.3** | 218 |
| Sapiens | 0.57 | 0.69 | 21.3 | 17.8 | **3** |
| Humatch (ours) | **0.77** | **0.82** | 20.6 | 13.0 | *33* |
| Experimental | 1.00 | 1.00 | 26.0 | 19.2 | - |

Figure 3 shows Humatch’s heavy and light chain mutation profiles closely match those from experiments. Vernier zone and interface residues that may affect the stability of the antibody are highlighted in orange and blue, respectively. Humatch made similar proportions of mutations within these zones (19.3% and 12.6% for the heavy and light chains, respectively) compared to those made experimentally (17.9% and 14.6%).

**Figure 3:**
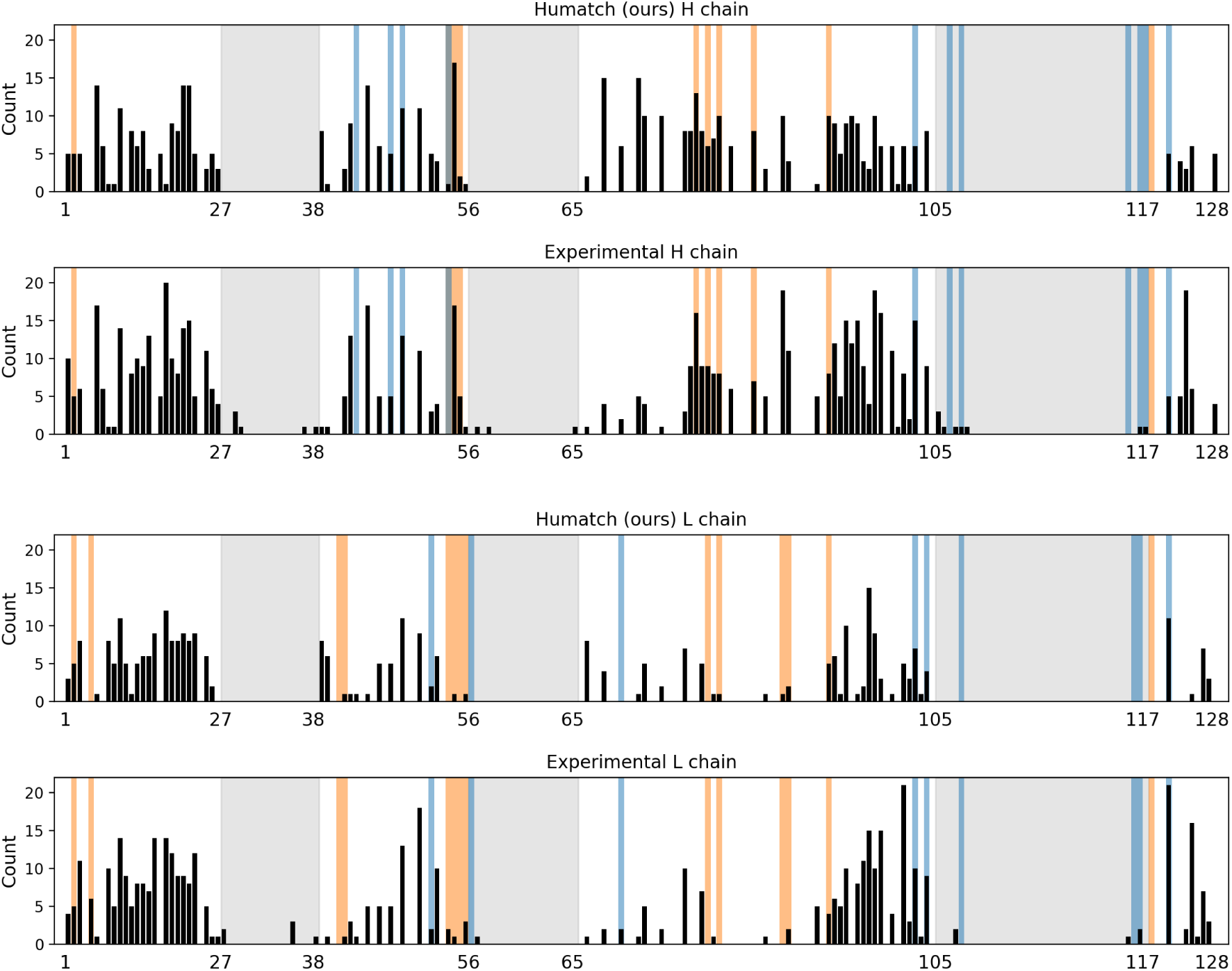
A comparison of Humatch and experimental mutation profiles for 25 precursor therapeutics, separated by heavy and light chains. The black bars show the number of mutations made at each IMGT sequence position across the 25 therapeutics. CDR regions are shaded in grey, Vernier zone residues in orange, and interface residues in blue.

Finally, we observe that when many mutations are made experimentally, Humatch also suggests more mutations and vice versa, with a Pearson correlation of 0.89. This correlation holds when considering the total number of heavy and light mutations and when considering each chain individually (see SI).

### 3.4 Humatch provides gene-specific humanisation while maintaining good heavy and light pairing

Humatch’s architecture and humanisation logic designs sequences with high experimental alignment (Table 2). This alignment is achieved while pushing designs towards well-paired specific V-genes and away from other human V-genes and non-human sequences (see Figure 2 and Methods). Here, we show that other humanisation tools optimise towards different V-genes than those chosen experimentally and that as none consider good VH/VL pairing this can lead to designs that Humatch predicts are unlikely to pair naturally.

As described above, we found that high CNN-P scores correlate with greater thermal stability (see SI). High CNN-P scores are therefore desirable for stable therapeutic designs. Figure 4 (top right) shows how Humatch’s CNN-P ranks all 25 computational and experimental designs summarised in Table 2 for their VH/VL pairing.

**Figure 4:**
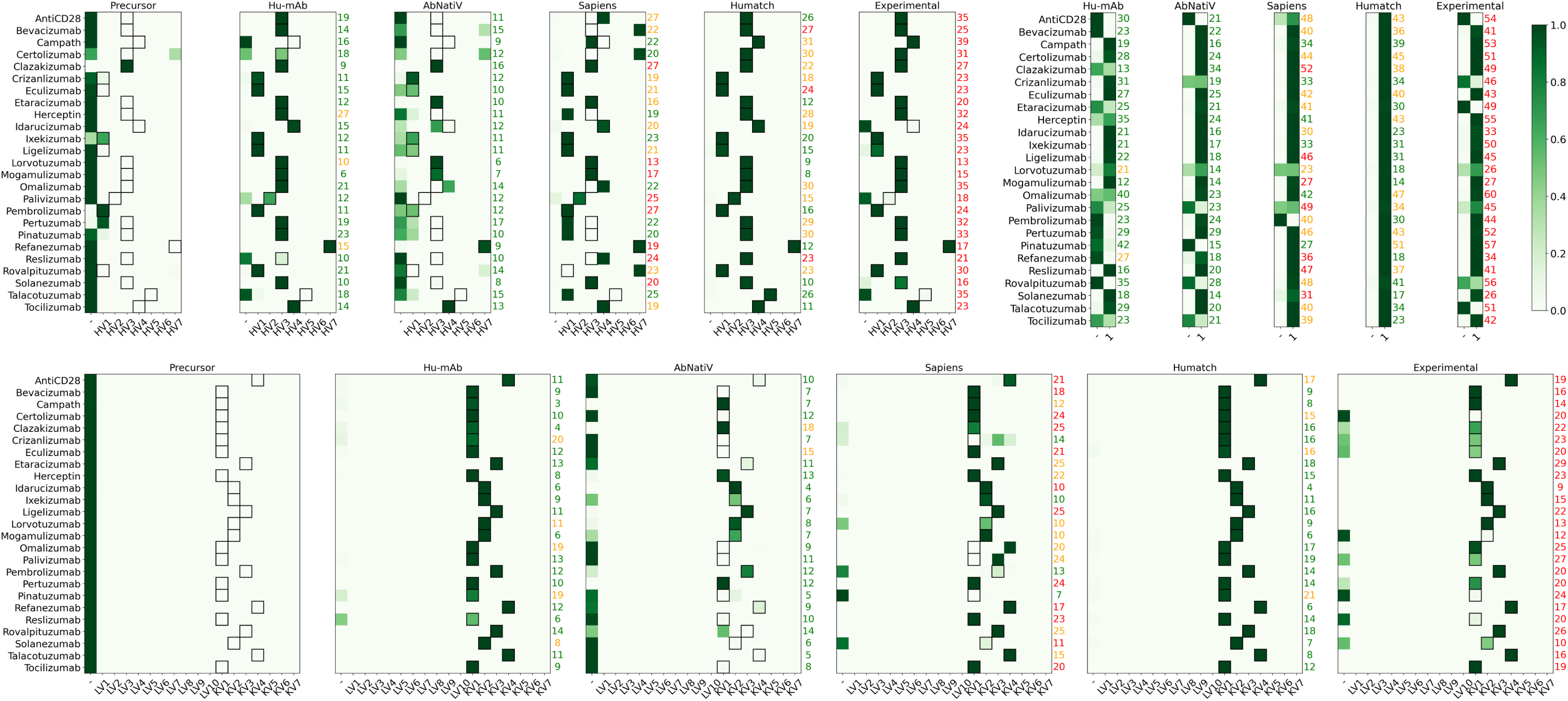
Comparison of Humatch scores achieved by different tools when humanising 25 precursor therapeutics. Humatch outputs multiclass prediction vectors for the heavy, light, and paired models. The predicted values are coloured according to the colour bar, with higher predictions shown in darker green. The target genes, shown as black boxes, were determined from the experimentally humanised sequences. The black boxes are omitted from the paired model for clarity - in this instance, we target class ‘1’, corresponding to the heavy and light chain being well paired. The edit distances of each design are shown to the right of each plot. Edit distances are coloured according to their relative size compared to experimental edits. Designs with the same or more edits as the experimental are coloured red, designs with fewer than 75% of the number of experimental edits are shown in green, and designs with edits between 75% and 100% of the experimental are shown in orange.

Hu-mAb’s designs are predicted to have the least natural pairing. Some designs from Sapiens and AbNatiV are also predicted to be ill-matched, as are several experimental endpoints (unfortunately, we could only find stability data for 13 of the 20 experimental endpoints predicted to pair well and none for the five predicted to pair less well for validation). The edit distances from each precursor sequence are shown to the right of each plot. Humatch designs VH/VL sequences that are predicted to be well-matched, according to our CNN-P, in fewer edits than were made experimentally for all 25 therapeutics. A lack of ground truth ‘badly’ paired sequence data for training CNN-P could result in both false positive and negative predictions; nevertheless, Humatch offers an improvement over ignoring VH/VL pairing concerns during humanisation.

Figure 4 (top left and bottom) highlights how other humanisation tools - AbNativ and Sapiens - often target different heavy and light V-genes than the experimentally designed endpoints. Outputs from these tools could also theoretically lie ‘between’ genes given their humanness scoring protocols. We find no direct evidence of this for these 25 designs (see Figure 4) but in the SI we show that Humatch successfully ranks more artificial ‘mixed-gene’ designs as non-human (51%) than AbNatiV (28%), Hu-mAb (8%), and OASis (1%).

Despite humanising towards different genes, most designs from Sapiens are still ranked as human by Humatch’s CNN-H and CNN-L classifiers. Fewer designs from AbNatiV are ranked as human by Humatch. Disagreement between humanisation tools is common as each may identify different patterns in the training data, even when trained on the same or similar datasets. Like Humatch, other tools also rank some experimentally humanised sequences low - Marks *et al.* find Hu-mAb ranks only ~66% of experimentally humanised VH/VL pairs as human.

Most of Hu-mAb’s designs achieve high CNN-H and CNN-L scores and target the same genes as experiments. In some cases, Hu-mAb, and other tools, achieve these high scores in fewer edits than our Humatch protocol. However, in many of these instances, these designs do not meet the target Humatch score given by the experimental sequence. As mentioned previously, Hu-mAb’s designs are also predicted to be less well-paired. If desired, Humatch can design sequences with high CNN-H, CNN-L, and CNN-P scores in fewer edits than stated in Figure 4 at the expense of lower germline-likenesses (see SI).

In summary, Figure 4 illustrates how Humatch is the only humanisation tool that allows both humanisation towards specific V-genes and good VH/VL pairing.

### 3.5 Humatch humanises sequences rapidly, allowing for high throughput computational design

To test Humatch in a high-throughput environment, we randomly selected 1,000 non-human naturally paired sequences from OAS as potential precursor therapeutics. Humatch scored these starting sequences and the top-scoring heavy and light genes were chosen as target genes during humanisation. Similarly to before, heavy, light, and paired target thresholds were set at 0.95 and the humanisation of each pair of sequences was allowed to continue up to a maximum combined edit distance of 60 from the starting sequence.

As Humatch includes logic to avoid local minima (see Figure 2 and Methods) it successfully humanised most sequences according to the above criteria, though 12 VH/VL pairs failed to humanise after hitting the maximum edit distance threshold. The mean heavy and light edit distances of the humanised sequences from their precursors were 11.6 and 13.9, respectively. These edit distances are lower than those reported for the 25 therapeutics earlier (Table 2), reflecting the higher experimental target thresholds used in some cases. The mean time required for humanisation was 36s when run on a standard desktop computer with 16 CPUs.

## 4 Discussion

Humanisation is a critical step in the design of many antibody therapeutics[3]. However, experimental optimisation involves trial and error and is time and cost-intensive[1]. Computational tools exist to aid in this process, though many are limited by long runtimes, no option to specify target germlines, and a lack of coherent joint heavy and light chain optimisation[10, 13, 17].

To support the rapid design of low immunogenicity, humanised antibody therapeutics, we have developed Humatch. Humatch is a collection of three CNN classification models that accurately identify human heavy sequences (CNN-H), human light sequences (CNN-L), and well-paired antibody VH/VL pairs (CNN-P). Humatch uses these classifiers and gene-specific germline data to guide its humanisation process through an iterative cycle of designing and ranking all possible single-point variants.

Humatch’s heavy and light antibody V-gene classifiers offer wider applications than previous methods as they are trained on a more extensive set of germlines and non-human species, including both human LV9 and KV7 genes and rhesus sequences. Humatch achieves near-perfect classification accuracies for most genes, though we avoid targeting the absolute highest accuracies to prevent overfitting, and instead attempt to favour a smoother humanisation process.

To further smooth the humanisation process, sequences are encoded using Kidera feature vectors. One-hot encodings were trialled and provided similar classification accuracies but designed sequences with lower experimental overlap. We hypothesise that due to their sparse nature and the fact that each convolutional filter acted over the entire sequence, one-hot features allowed ‘nonsense’ mutations never observed in human or non-human sequences to receive high CNN predictions by chance. Kidera feature vectors provided near-perfect classification accuracies so were selected over more descriptive Large Language Model (LLM) embeddings to restrict the size of Humatch’s model weights and runtime.

Humatch’s CNN-H and CNN-L classifiers output multiclass predictions (e.g. non-human plus all human heavy V-genes), meaning that during humanisation they push sequences not only towards their target gene but also away from other human genes. Gene-specific amino acid frequency lookup tables, similar to chain-level PSSMs used by AbNatiV, also help in this guidance, avoiding humanising sequences ‘between’ genes. Mixed-gene sequences are not observed naturally and so pose potential immunogenic risks, yet we show many are ranked highly by other humanisation tools.

With its CNN-P, Humatch also ensures heavy and light chains remain well paired during the humanisation process. This offers potential expression and thermostability benefits as high CNN-P predictions are shown to correlate with higher melting temperatures. CNN-P also considers the fact that immunogenic epitopes could be formed between heavy and light chains - therapeutics with the highest anti-drug antibody levels have low CNN-P predictions (see SI). Some caution should be applied when interpreting these predictions for other tools though given the lack of 100% ground truth ‘badly’ paired data in OAS. Nevertheless, Humatch is the only humanisation tool that explicitly optimises for well-matched sequences.

Generally, it is difficult to compare the classification and humanisation outputs of different humanisation tools. Even when trained on the same or similar data, various architectures can learn to identify and prioritise different patterns in the data. For this reason, we prioritised high experimental overlap and germline-likeness in optimising Humatch’s architecture and humanisation protocol. Humatch’s designs overlap with experimental designs more than any other method - 77% and 82% of heavy and light mutations match exactly. This high overlap is achieved despite Humatch making the second-largest number of mutations of all methods, indicating that Humatch does not simply find the ‘easy’ mutations.

Finally, Humatch offers fast humanisation by targeting an initial germline-likeness score that is calculated based on observed therapeutics. This initial step takes less than a second to complete, immediately sets Humatch on a sensible humanisation trajectory to any target gene, and allows for high throughput experiments with a total runtime of ~35s per paired VH/VL. Given Humatch’s quick runtime, strong experimental alignment, and inbuilt guidance towards single germlines and natural chain pairings, it should speed up and reduce the cost of the drug discovery process.

## 5 Methods

### 5.1 Classifier training data preparation

Humatch was trained on data from the Observed Antibody Space (OAS)[11, 12] database. Both unpaired and paired sequences were processed to remove redundancy. Sequences lacking Cystines at IMGT positions 23 and 104 were also removed. Finally, we required that both the first (IMGT position 1) and last (IMGT positions 128 and 127 for the heavy and light chains respectively) residues were present. As Humatch has been trained on complete VH and VL sequences, users should only input complete sequences for reliable results.

Table 3 shows the total number of heavy and light sequences used to train Humatch’s unpaired CNNs, broken down by V-gene. We did not split our data by D- or J-genes, so each V-gene class includes a mix of these other genes. In total, 8.26m human heavy sequences and 12.73m human light sequences were used for training, alongside 3.77m and 1.41m non-human heavy and light sequences, respectively. This data was subject to a stratified 80-10-10 train-validation-test split.

**Table 3:** Total number of human sequences used to train, validate and test Humatch’s heavy and light chain CNN classifiers, broken down by V-gene. Heavy and Kappa V-genes 8-10 do not exist in OAS. Lambda and Kappa sequences were used together to train CNN-L.

|  | Heavy | Lambda | Kappa |
| --- | --- | --- | --- |
| V1 | 2,107,242 | 1,100,881 | 3,073,544 |
| V2 | 263,277 | 2,266,529 | 974,970 |
| V3 | 2,949,663 | 1,112,650 | 2,878,296 |
| V4 | 2,436,892 | 72,589 | 867,144 |
| V5 | 386,538 | 27,716 | 9,065 |
| V6 | 35,566 | 107,905 | 46,063 |
| V7 | 84,216 | 114,900 | 4,122 |
| V8 | - | 36,861 | - |
| V9 | - | 21,353 | - |
| V10 | - | 13,678 | - |

Humatch’s paired CNN was trained on 1,673,734 paired human sequences from OAS, and 5,009,443 artificially ‘badly’ paired human sequences. Both datasets were processed and split similarly to our unpaired data. To increase the chances of our artificially paired sequences being poor pairs, we paired only unpaired heavy and light chains from different B-cell classes (e.g. memory vs naive) and different OAS studies.

IMGT numbering for all sequences was taken from OAS (originally numbered using AN-ARCI[22]). Sequences we padded and aligned to the 200 most common sequence positions identified by KASearch[19], more than the 152 positions used previously by Hu-mAb[10]. IMGT positions not present in a sequence were represented with pad tokens. Paired heavy and light chains were joined with a length-ten pad, creating a length 410 sequence. Sequences were encoded using Kidera feature vectors[20] and pad tokens were represented with zero vectors.

### 5.2 Humatch Convolutional Neural Network classifiers

Humatch consists of three CNN classifiers - one to identify human heavy V-genes (CNN-H), one for human light V-genes (CNN-L), and one for well-paired human sequences (CNN-P).

All three of Humatch’s CNNs included 40 convolutional filters, each with a kernel size of ten and a stride of one. The models’ inputs were encoded with ten-dimensional Kidera feature vectors and padded to be of size (batch size *×* sequence length *×* 10). The sequence length was 200 for unpaired sequences and 410 for paired sequences. When passed through the convolutional filters, the input was padded with zeroes at the start and end to ensure the filter outputs were the same length as the input. ReLU activations were used throughout.

Dropout layers with probabilities of 20% were applied following each convolutional filter. These outputs were then max-pooled with a pool size of two and a stride of one. Padding was not applied at this stage, so the lengths of the outputs were one less than the inputs. These outputs were then flattened.

Probabilities of the heavy and light chains being human and well-matched were obtained by passing the output above through two dense layers. These dense layers reduced the output dimensions first to 300 and then to the number of label classes for each CNN. Humatch’s heavy CNN classifier has an output dimension of eight (one non-human class and seven heavy V-genes), the light CNN has an out dimension of 18 (one non-human class, ten lambda V-genes, and seven kappa V-genes), and the paired CNN has an output dimension of two (fake and true pairs). A sigmoid activation function is applied to the final layer to ensure predictions across all classes sum to one.

Training of all models used Adam optimisation, binary cross-entropy loss, a learning rate of 0.000075, and a batch size of 1,024. Each model was trained for a maximum of 15 epochs. Weights were saved following each epoch and optimum weights were selected to ensure both high classification accuracy and smooth humanisation. This selection was manual - validation set precision and recall values were examined and example sequences were humanised using different combinations of CNN-H, CNN-L, and CNN-P weights.

Humatch’s CNN and training code used Python v3.9, TensorFlow v2.16, and Keras v3.0. Details of all dependencies used can be found at github.com/oxpig/Humatch.

### 5.3 Humatch humanisation logic

Humatch uses a combination of rapid germline lookups and CNN predictions to guide its humanisation process. All steps of this process are described below and summarised in Figure 2. If at any time the sequence meets all three CNN humanisation target thresholds (default 0.95 for the heavy, light, and paired CNNs) then the humanisation process stops. A maximum edit distance from the precursor sequence (default 60) can also be set to break the humanisation process early.

The humanisation process starts by calculating a ‘germline-likeness’ score for the input sequence. This score measures how much the sequence looks like the positive training data of the target gene (chosen by the user or selected automatically). Specifically, Humatch precomputes how frequently each of the 20 canonical amino acids is observed at every sequence position e.g. for HV1, ‘Q’ is found at IMGT position one 95.8% of the time, while ‘E’ is observed there 4.0% of the time. These frequencies are extracted for each residue in the input sequence and the mean is taken (padded positions receive zero scores). This germline-likeness score is then compared to a target value (default 0.40) calculated based on observed therapeutics (see SI). If the current score is below the target, mutations are made based on whichever will lead to the single largest increase in the germline-likeness score. This process happens separately for the heavy and light chains and continues until the thresholds for each chain are met. No mutations are made at this stage if a chain’s germline-likeness score already meets the target threshold.

After this initial quick germline shift of the input sequence, Humatch makes every possible single-point mutation (ignoring padded positions) and scores these with its three CNNs. Heavy chain mutations will affect the predictions of CNN-H and CNN-P only, while light chain mutations will impact CNN-L and CNN-P. CDR mutations are excluded by default in this step (this reduces the likelihood of disrupting binding and speeds up humanisation) though users can add or remove residues from consideration as desired. Once predictions have been made, net predictions are calculated for each CNN separately by subtracting the scores achieved by the unmutated sequence (the output of the germline-likeness step and Humatch’s current ‘best’ design).

The net predictions for each CNN are then scaled using the same germline-specific frequencies from step one. Each positive net prediction is multiplied by the observed frequency of the single-point mutation made. Negative net predictions are multiplied by one minus the frequency. The effect of this step is to prioritise predictions that both increase humanness (separately for each CNN) and are frequently observed. We scale the net predictions rather than the absolute probabilities so that mutations that improve a CNN score always rank above those that worsen it. Noise can be added at this stage by adding a fixed number to each scaling factor before scaling. Adding more noise reduces how much the humanisation process favours previously observed mutations. By default, Humatch uses low noise (0.01).

Once scaled, the net predictions are then made positive again by subtracting the lowest negative score across all three models from all adjusted net predictions. Choosing to add this fixed value, instead of re-adding the predictions of the best sequence, equalises the importance of all three CNN scores ahead of the next stage.

These positive predictions are then scaled by how far away each current CNN score is from its respective target before they are combined e.g. if the current sequence has a heavy CNN score of 0.10, a light CNN score of 0.97, and paired CNN score of 0.55, with targets of 0.95 for all three models, then the heavy predictions would be multiplied by 0.85, the light by zero (as the threshold is already met), and the paired by 0.40. The CNN-H and CNN-L predictions are then concatenated and added element-wise to the CNN-P predictions. This second scaling stage before summing ensures Humatch does not waste mutations optimising one trait beyond the target threshold while others remain low.

Once summed, the top-ranked variant (that has not been selected in previous rounds) is chosen as Humatch’s new best sequence. If this new sequence surpasses all three CNN thresholds or the maximum allowed edit distance is reached, the humanisation process stops. Otherwise, the humanisation process, excluding the initial rapid germline-likeness matching, repeats.

Note that in the above example, as CNN-P considers both heavy and light mutations, a light chain mutation could still be selected despite the CNN-L threshold being met. In this instance, the CNN-L score of the newly selected sequence could differ from the previous best sequence (positively or negatively). If the light chain CNN score were to drop back below the target threshold following this, then the CNN-L scores would be considered again in the next round of mutations.

## Funding

This work was supported by the Biotechnology and Biological Sciences Research Council (BB-SRC) and GlaxoSmithKline (GSK).

### Conflict of Interest

None declared.

## Supporting information

Humatch Supplementary Information

## Notes

https://github.com/oxpig/Humatch

https://zenodo.org/records/13764770

### Competing Interest Statement

The authors have declared no competing interest.

