## Supplementary material for "Humatch - fast, gene-specific joint humanisation of antibody heavy and light chains": Humatch Supplementary Information

September 16, 2024

#### 1 Humatch classification accuracy when trained and tested on Hu-mAb's data

In the main text, we highlight how Humatch was trained on a more extensive dataset than used by Hu-mAb[1], and that training is stopped early to facilitate smoother humanisation. Table S1 demonstrates that Humatch's architecture is capable of achieving comparably high accuracies to Hu-mAb when trained and tested on the same dataset. Here, Humatch's CNN-H and CNN-L were trained for three epochs, similarly to the main text, but this time on Hu-mAb's data. The data consists of only 152 IMGT residue positions, the light chain data lacks V-genes LV9 and KV7, and neither heavy nor light datasets contain rhesus sequences in the negative training data. Table S1 shows Humatch achieved perfect PR AUC and ROC AUC scores for all V-genes measured to three decimal places.

| Class | PR AUC | F1 | MCC | ROC AUC | BA |
|---|---|---|---|---|---|
| HV1 | 1.000 | 1.000 | 1.000 | 1.000 | 1.000 |
| HV2 | 1.000 | 1.000 | 1.000 | 1.000 | 1.000 |
| HV3 | 1.000 | 1.000 | 1.000 | 1.000 | 1.000 |
| HV4 | 1.000 | 1.000 | 1.000 | 1.000 | 1.000 |
| HV5 | 1.000 | 1.000 | 1.000 | 1.000 | 1.000 |
| HV6 | 1.000 | 1.000 | 1.000 | 1.000 | 1.000 |
| HV7 | 1.000 | 1.000 | 1.000 | 1.000 | 1.000 |
| LV1 | 1.000 | 1.000 | 1.000 | 1.000 | 1.000 |
| LV2 | 1.000 | 1.000 | 1.000 | 1.000 | 1.000 |
| LV3 | 1.000 | 1.000 | 1.000 | 1.000 | 1.000 |
| LV4 | 1.000 | 1.000 | 1.000 | 1.000 | 1.000 |
| LV5 | 1.000 | 1.000 | 1.000 | 1.000 | 1.000 |
| LV6 | 1.000 | 1.000 | 1.000 | 1.000 | 1.000 |
| LV7 | 1.000 | 1.000 | 1.000 | 1.000 | 1.000 |
| LV8 | 1.000 | 1.000 | 1.000 | 1.000 | 1.000 |
| LV10 | 1.000 | 0.998 | 0.998 | 1.000 | 1.000 |
| KV1 | 1.000 | 1.000 | 1.000 | 1.000 | 1.000 |
| KV2 | 1.000 | 1.000 | 1.000 | 1.000 | 1.000 |
| KV3 | 1.000 | 1.000 | 1.000 | 1.000 | 1.000 |
| KV4 | 1.000 | 1.000 | 1.000 | 1.000 | 1.000 |
| KV5 | 1.000 | 1.000 | 1.000 | 1.000 | 1.000 |
| KV6 | 1.000 | 1.000 | 1.000 | 1.000 | 1.000 |

Table S1: Performance of Humatch's heavy and light CNN classifiers trained and tested on HumAbs dataset. Sequences belonging to all genes are classified with near-perfect accuracy. PR AUC = Area Under the Precision-Recall Curve; F1 = F1-score, MCC = Matthews Correlation Coefficient; ROC AUC = Area Under the Receiver Operating Characteristic Curve; BA = Balanced Accuracy.

### 2 Heavy-light gene pairings of Humatch's true and artificially paired training data

The artificially paired data used to train Humatch must have approximately the same heavy-light gene pairings as the true data so that CNN-P cannot simply learn to identify unusual pairings. Figure S1 shows that both datasets have similar distributions of pairings, with most pairs belonging to heavy V-genes 1, 3, and 4. Most light genes belong to Kappa genes 1-4 and Lambda genes 1-3.

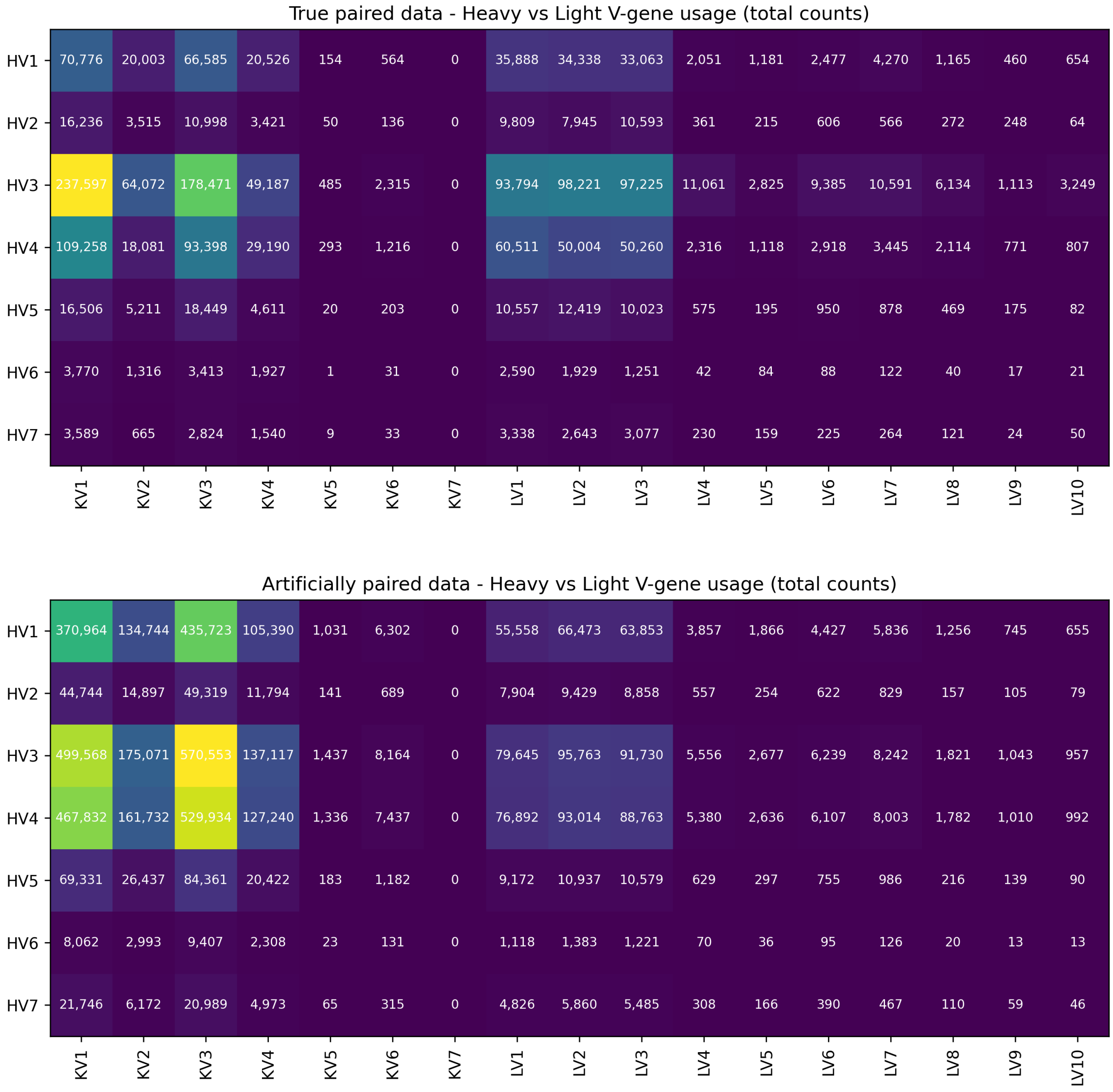


Figure S1: Distribution of heavy-light gene pairings for the true (top) and artificially 'badly' paired (bottom) datasets used to train Humatch. The total counts for each gene pair are given in the respective cells and these are coloured by their proportion of the total dataset (low proportions are shown in purple and high proportions in yellow).

### 3 Humatch's anti-drug antibody correlation

Figure S2 shows how each CNN contributes to screening for therapeutics that may illicit anti-drug antibody (ADA) responses in high proportions of patients. All CNNs offer some separation of low- and high-ADA antibodies, though CNN-H and CNN-L both include one high-ADA therapeutic in their top-ranked bin, unlike CNN-P.

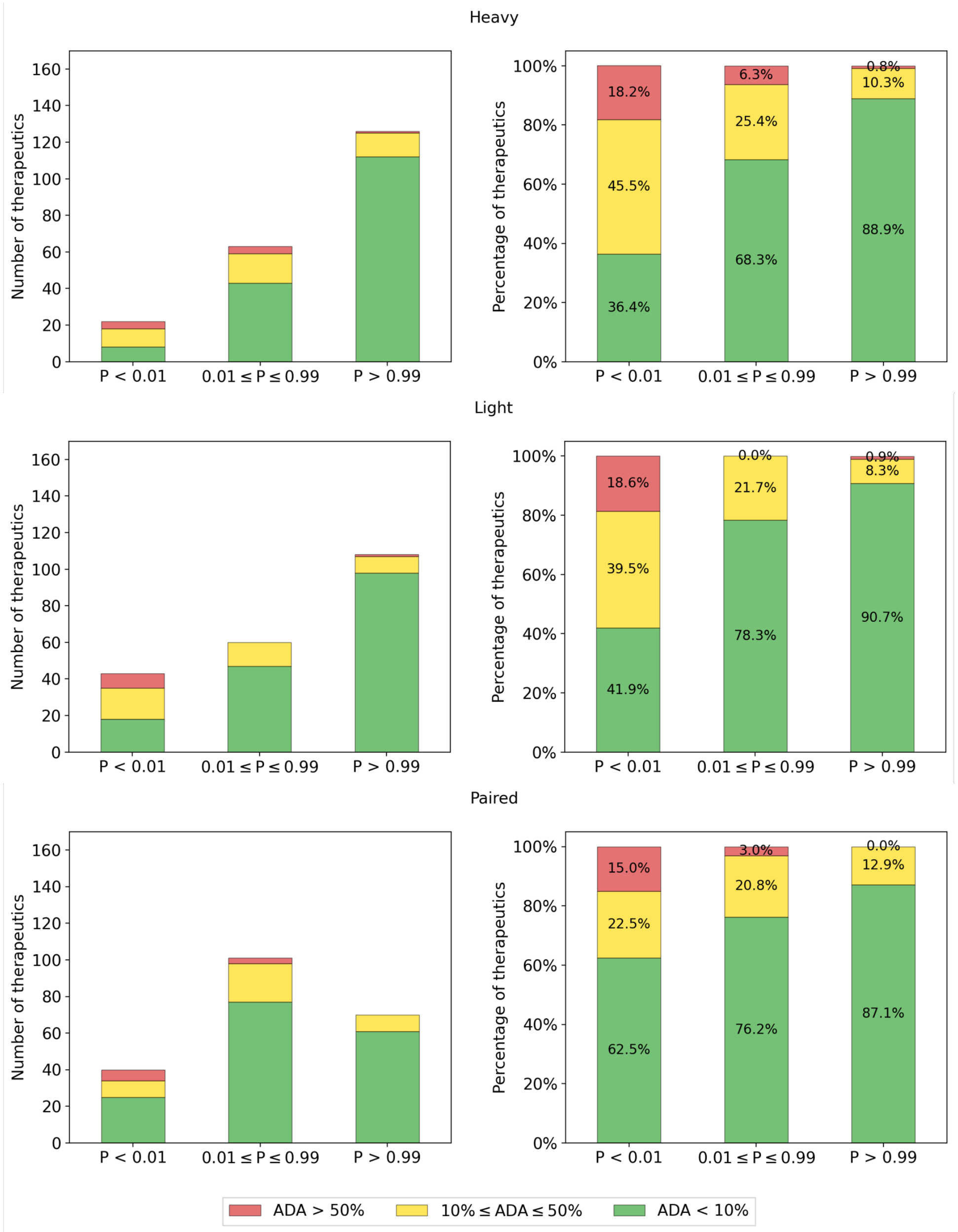


Figure S2: Comparison of how each CNN separates 211 therapeutics with different anti-drug antibody (ADA) levels.

In the main text, we highlight how the minimum of the three scores is recommended to remove most high ADA therapeutics. However, Figure S3 shows that the maximum and mean of the three scores are also efficient at screening out only highly immunogenic therapeutics while maintaining most others.

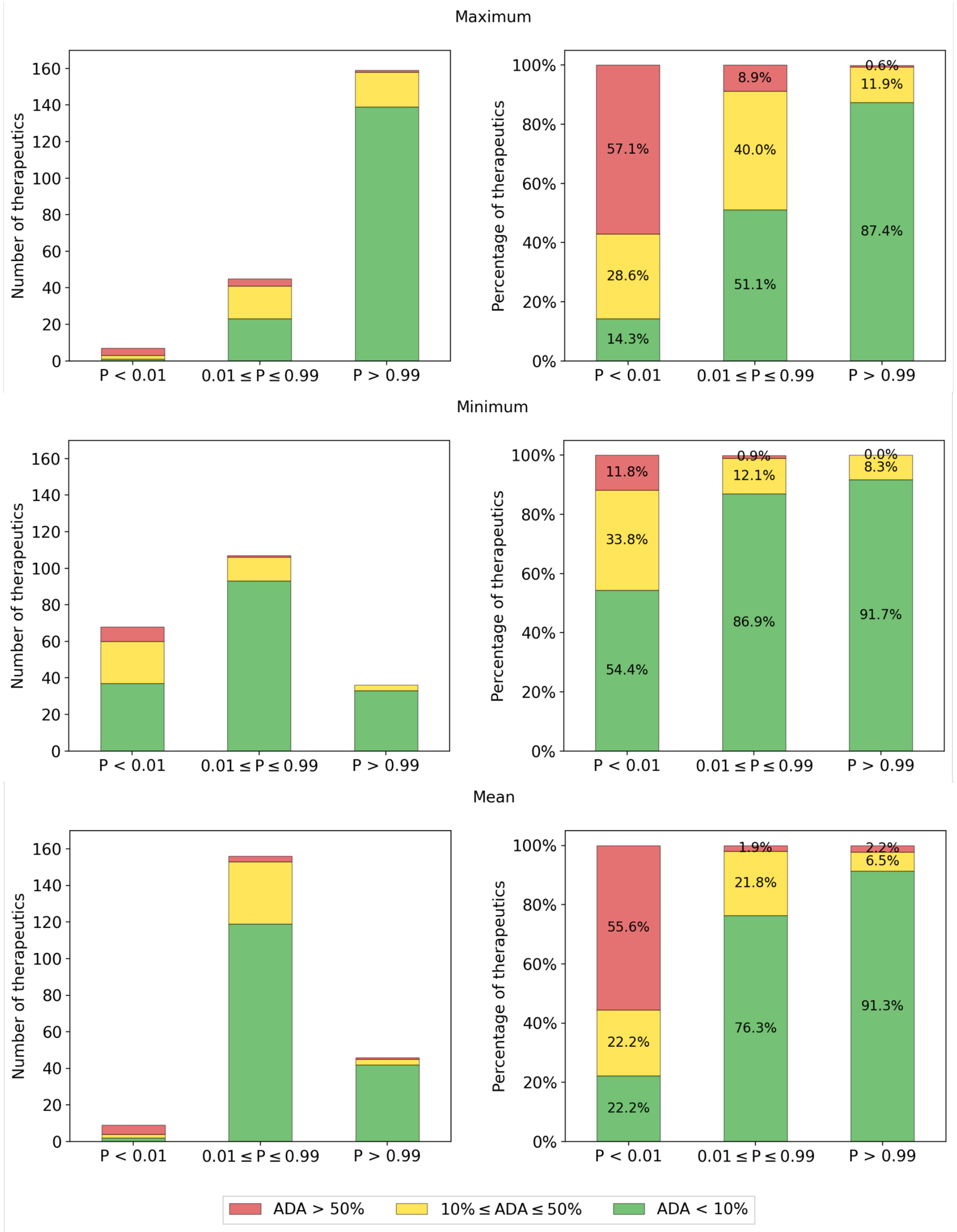


Figure S3: Comparison of how the maximum, minimum, or mean of the three CNNs separates 211 therapeutics with different anti-drug antibody (ADA) levels.

### 4 Hu-mAb, AbNatiV, and OASis' anti-drug antibody correlations

Hu-mAb[1], AbNatiV[2], and OASis[3] were used to score the same 211 therapeutics with ADA information as Humatch. These tools were run with default and paper-recommended parameters.

Similarly to Humatch, the minimum of each tool's heavy and light chain scores were taken and used to group the 211 therapeutics into three bins (Figure S4). The bin thresholds were selected to yield approximately the same number of therapeutics in each bin as Humatch.

We found that all methods placed the same number of high ADA (red) therapeutics in the bottom bin. Hu-mAb placed proportionally more medium (yellow) ADA therapeutics in this bin than other methods. Hu-mAb and OASis also achieved cleaner top bins than Humatch and AbNatiV, with all (or all but one) therapeutics having low (green) ADA levels.

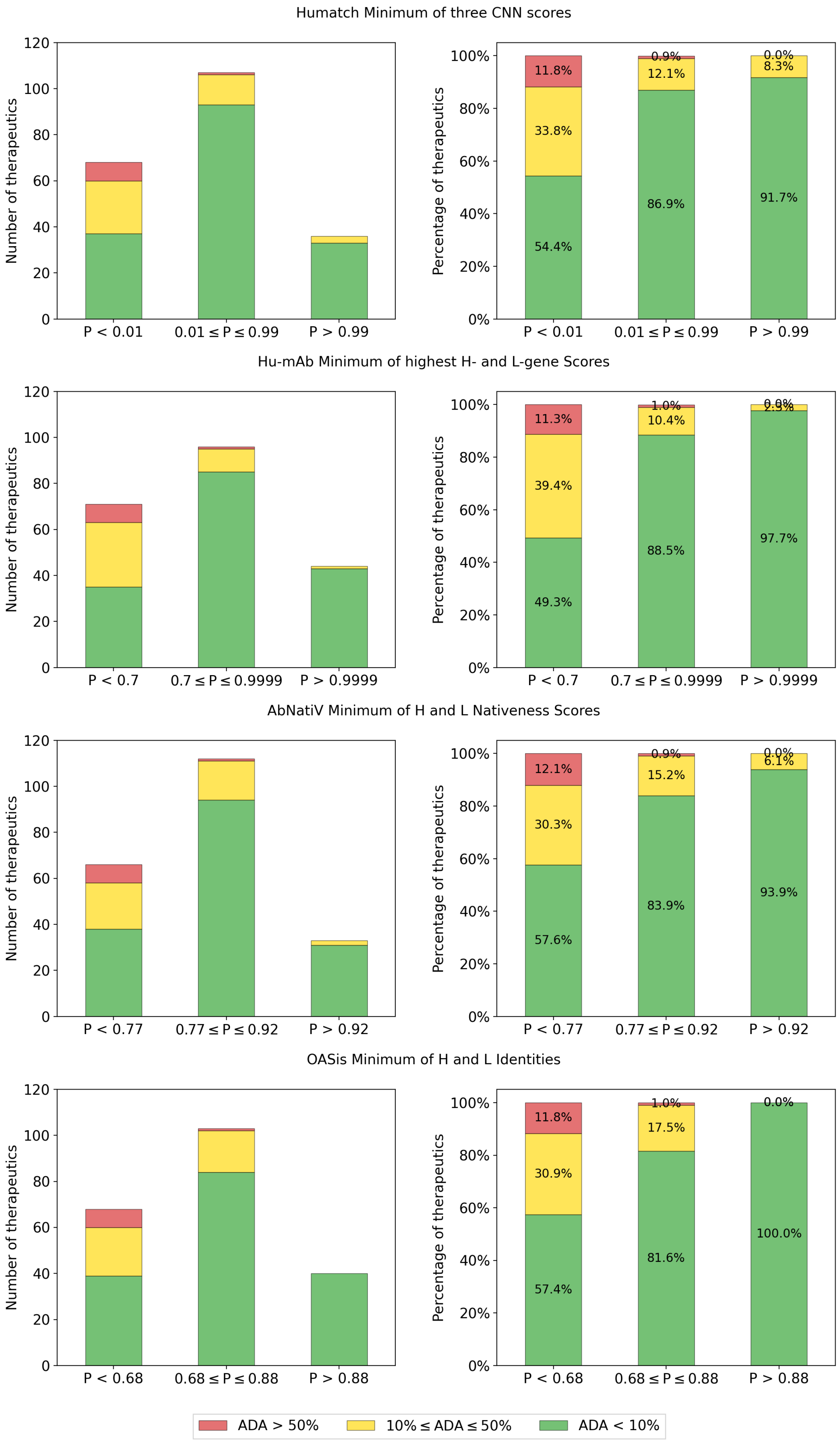


Figure S4: Comparison of how Humatch, Hu-mAb, AbNatiV, and OASis separate 211 therapeutics with different anti-drug antibody (ADA) levels.

### 5 CNN-P thermostability correlation

To evaluate whether CNN-P can identify better expressing or more stable antibodies, we used it to score 137 therapeutics from Jain *et al.*[4] (Figure S5). In total, 98 therapeutics received high scores (CNN-P $> 0.5$) and 39 had low scores (CNN-P $\leq 0.5$). While expression, measured using HEK titers, showed no significant correlation according to a two-sided T-test ($p = 0.13$), higher CNN-P scores were shown to correlate with greater melting temperatures ($p = 0.00092$).

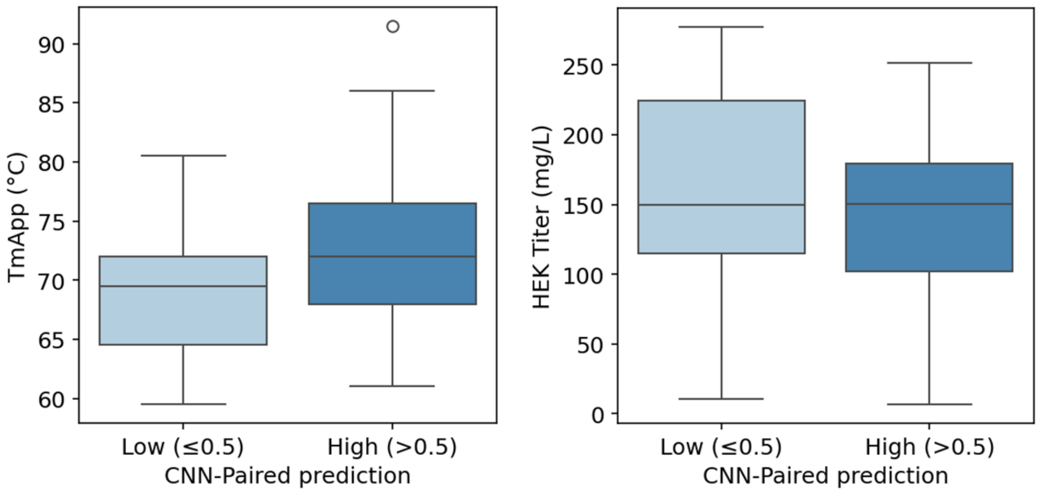


Figure S5: Box plots showing the melting temperatures (left) and expression levels (right) of 137 therapeutics from Jain *et al.* grouped by Humatch's CNN-P scores.

### 6 Choosing optimum CNN classifier cutoffs

The data used for training Humatch suffers from large gene-class imbalances. Therefore, to determine optimum classifier cut-offs, we used sample weights to counter this imbalance. We recommend using a threshold of 0.95 for all classifiers as this was found to provide high sample-weighted precision and recall for all classes (Table S2). We prioritised near-perfect precision over recall in selecting this threshold given the nature of the humanisation problem.

| Class | Precision | Recall |
|---|---|---|
| HV1 | 0.999 | 0.946 |
| HV2 | 1.000 | 0.980 |
| HV3 | 0.999 | 0.945 |
| HV4 | 1.000 | 0.959 |
| HV5 | 1.000 | 0.975 |
| HV6 | 1.000 | 0.780 |
| HV7 | 1.000 | 0.913 |
| LV1 | 1.000 | 0.998 |
| LV2 | 1.000 | 0.994 |
| LV3 | 1.000 | 0.997 |
| LV4 | 1.000 | 1.000 |
| LV5 | 1.000 | 0.980 |
| LV6 | 1.000 | 1.000 |
| LV7 | 1.000 | 0.981 |
| LV8 | 1.000 | 1.000 |
| LV9 | 1.000 | 0.995 |
| LV10 | 1.000 | 0.969 |
| KV1 | 1.000 | 0.973 |
| KV2 | 1.000 | 0.988 |
| KV3 | 1.000 | 0.989 |
| KV4 | 1.000 | 0.997 |
| KV5 | 1.000 | 0.998 |
| KV6 | 1.000 | 0.993 |
| KV7 | 1.000 | 0.295 |
| true pairs | 0.995 | 0.933 |

Table S2: Comparison of the sample-weighted precision and recall achieved by each class using classifier cut-offs of 0.95. The sample-weighting used equalised the class imbalance between the target class and all other classes in each instance. The precision exceeds 0.999 for all CNN classes. The recall was high for most classes, though dropped when little training data was available e.g. KV7.

### 7 Humatch classification accuracy remains high when data is split by allele

To test whether Humatch's predictions might be robust to classifying new human alleles yet to be recorded, we split the training data so that the test set contained only alleles unseen during training for heavy V-genes 1-4. We selected only heavy V-genes 1-4 for this task as others did not allow us to obtain sufficient data to maintain the same train-validation-test dataset sizes and proportions as used in Humatch's full training.

CNN-H was therefore retrained with the same negative data and heavy V-genes 5-7 as the main text. HV1 alleles 1-11, HV2 alleles 1-4, HV3 alleles 1-4, and HV4 alleles 1-7 were also used for training and validation. Other HV1-4 alleles were reserved for testing, alongside the same negative and HV5-7 data as before. The training was stopped after three epochs, similar to the main text.

Tables S3 & S4 show that Humatch's test set performance remains high for all genes, regardless of whether or not the data was split by allele. The largest drop in performance was for HV2 which may be because the amount of training data available is an order of magnitude lower than HV1, 3, and 4.

| Class | PR AUC | F1 | MCC | ROC AUC | BA |
|---|---|---|---|---|---|
| HV1 | 1.000 | 0.998 | 0.998 | 1.000 | 1.000 |
| HV2 | 0.949 | 0.909 | 0.911 | 0.998 | 0.918 |
| HV3 | 0.995 | 0.952 | 0.936 | 0.998 | 0.975 |
| HV4 | 1.000 | 0.990 | 0.987 | 1.000 | 0.996 |
| HV5 | 1.000 | 0.987 | 0.987 | 1.000 | 1.000 |
| HV6 | 0.991 | 0.811 | 0.825 | 1.000 | 0.999 |
| HV7 | 0.991 | 0.914 | 0.915 | 1.000 | 0.987 |

Table S3: Performance of Humatch's heavy CNN classifier, CNN-H, when trained and tested on data split by allele (HV1-4 only). Sequences belonging to all genes are classified with high accuracy. Gene classes with the lowest scores, such as HV2, were tested on unseen alleles and have comparatively little training data available. PR AUC = Area Under the Precision-Recall Curve; F1 = F1-score, MCC = Matthews Correlation Coefficient; ROC AUC = Area Under the Receiver Operating Characteristic Curve; BA = Balanced Accuracy.

| Class | Precision | Recall |
|---|---|---|
| HV1 | 0.999 | 1.000 |
| HV2 | 1.000 | 0.820 |
| HV3 | 0.998 | 0.888 |
| HV4 | 0.999 | 0.986 |
| HV5 | 1.000 | 0.997 |
| HV6 | 1.000 | 0.909 |
| HV7 | 1.000 | 0.913 |

Table S4: Comparison of the sample-weighted precision and recall achieved by each gene using classifier cut-offs of 0.95 for CNN-H trained on allele-split data. The sample-weighting used equalised the class imbalance between the target class and all other classes in each instance. The precision exceeds 0.99 for all CNN classes. The recall was high for most classes, though dropped when little training data was available and the test set included previously unseen alleles e.g. HV2.

### 8 Germline-likeness of human, non-human, and therapeutic antibodies

Humatch targets an initial germline-likeness score of 0.40 during its humanisation process for both heavy and light chains. This cutoff was determined based on the lower 20th percentile of germline-likeness scores achieved by 744 approved and phase 1–3 therapeutics obtained from Thera-SAbDab (August 2024) and Marks *et al.*[1]. Figure S6 shows the distribution of these therapeutic scores (black) compared to a sample of non-human (red) and human (green and blue) sequences split by V-gene (lower number genes are darker). Non-human genes were scored against the highest-ranked human gene determined by Humatch. Some non-human heavy and light sequences already surpass the 0.40 threshold used. In these instances, these sequences would not be altered in the initial germline-likeness step of the humanisation process.

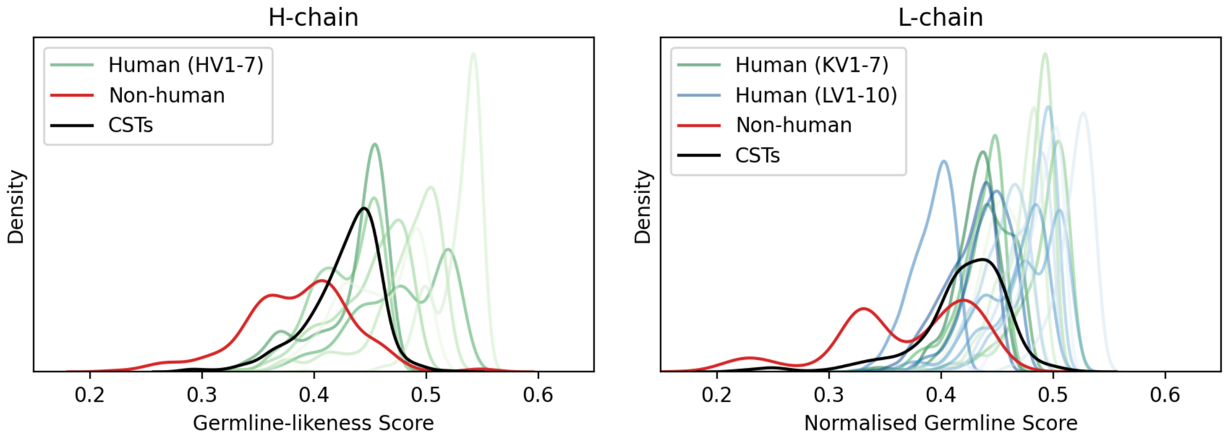


Figure S6: Comparison of the germline likeness scores for a random selection of heavy and light chain sequences from OAS and 744 clinical-stage therapeutics (CSTs). As expected, on average we observe that human sequences (green and blue, lower number genes are darker) have higher human germline-likeness scores than non-human sequences (red). Therapeutic sequences (black) tend to have high germline-likeness scores, comparable to human sequences.

### 9 Baseline humanisation of 25 therapeutics using top germline mutations only

We conducted a baseline experiment where only the top germline mutations were made to test the importance of Humatch's CNN guidance in the humanisation process. In this baseline, the same number of heavy and light chain mutations were made as in Humatch's full humanisation logic. We found that no baseline-designed paired sequences matched Humatch's designs exactly and only two designs passed all three CNN target thresholds of 0.95 (Table S7). This baseline shows that Humatch considers the sequence context when deciding which germline mutations should be made.

| Therapeutic | H-edit | L-edit | CNN-H | CNN-L | CNN-P |
|---|---|---|---|---|---|
| AntiCD28 | 8 | 10 | 1.00 | 1.00 | 0.00 |
| Bevacizumab | 6 | 6 | 0.71 | 0.00 | 1.00 |
| Campath | 10 | 4 | 0.00 | 1.00 | 1.00 |
| Certolizumab | 4 | 2 | 1.00 | 0.37 | 1.00 |
| Clazakizumab | 2 | 0 | 1.00 | 0.99 | 1.00 |
| Crizanlizumab | 6 | 4 | 0.99 | 0.05 | 0.00 |
| Eculizumab | 8 | 6 | 1.00 | 0.04 | 0.63 |
| Etaracizumab | 10 | 4 | 1.00 | 1.00 | 0.03 |
| Herceptin | 4 | 2 | 0.97 | 0.99 | 1.00 |
| Idarucizumab | 2 | 4 | 0.92 | 0.10 | 1.00 |
| Ixekizumab | 4 | 9 | 0.99 | 0.70 | 1.00 |
| Ligelizumab | 4 | 2 | 0.55 | 1.00 | 0.97 |
| Lorvotuzumab | 6 | 11 | 1.00 | 0.83 | 0.00 |
| Mogamulizumab | 4 | 6 | 0.53 | 0.03 | 0.99 |
| Omalizumab | 6 | 5 | 0.95 | 0.02 | 0.08 |
| Palivizumab | 10 | 8 | 0.00 | 1.00 | 0.25 |
| Pembrolizumab | 8 | 2 | 0.99 | 1.00 | 0.32 |
| Pertuzumab | 4 | 2 | 0.47 | 0.86 | 0.99 |
| Pinatuzumab | 4 | 6 | 1.00 | 0.10 | 0.98 |
| Refanezumab | 10 | 4 | 0.98 | 0.48 | 0.99 |
| Reslizumab | 4 | 4 | 1.00 | 0.01 | 0.97 |
| Rovalpituzumab | 5 | 2 | 0.99 | 1.00 | 0.55 |
| Solanezumab | 4 | 8 | 0.00 | 0.03 | 0.94 |
| Talacotuzumab | 10 | 8 | 0.46 | 0.73 | 1.00 |
| Tocilizumab | 4 | 6 | 0.12 | 0.00 | 1.00 |

Figure S7: CNN scores achieved by our baseline germline-only humanisation method. 'H-edit' and 'L-edit' describe the edit distances between Humatch's and baseline designs. CNN thresholds of 0.95 passed are shown in green. Lower scores are shown in red. Only 2/25 baseline designs passed all three thresholds.

### 10 Humatch humanisation of 25 precursor therapeutics

We observe that when many mutations are made experimentally, Humatch also suggests more mutations and vice versa. This relation holds for the number of heavy edits (Pearson $r = 0.86$), light edits ($r = 0.90$), and the total number of edits ($r = 0.89$) when tested on 25 precursor sequences from Marks *et al.* with known experimental endpoints (Figure S8).

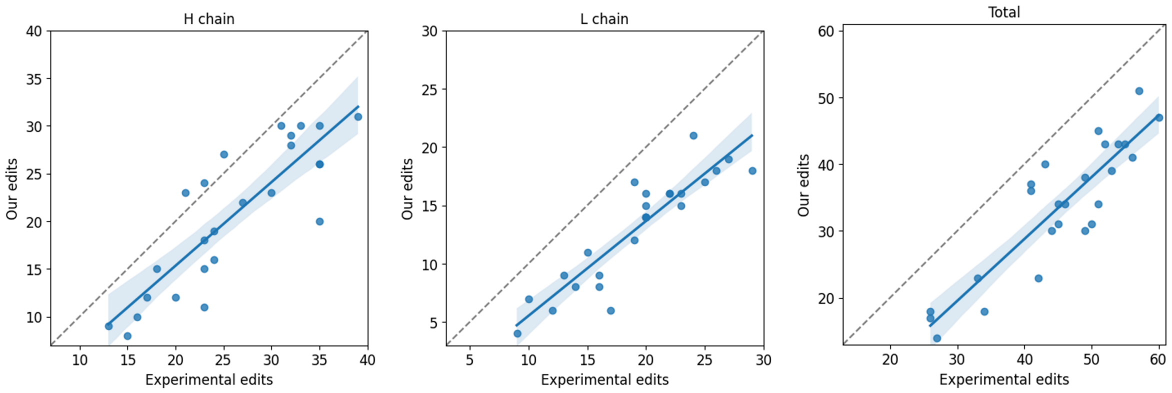


Figure S8: Comparison of the number of experimental edits made during the humanisation of 25 precursor therapeutics and the number suggested by Humatch. Pearson correlations of 0.86, 0.90, and 0.89 exist for the heavy, light, and total number of edits respectively.

### 11 Humatch humanisation of 25 precursor therapeutics using a lower initial germline-likeness

The initial germline-likeness (GL) targeted by Humatch can be adjusted by the user. By default, Humatch uses a target of 0.40, corresponding to the lower 20th percentile of GL scores achieved by clinical-stage therapeutics (CSTs). Lowering this target can allow Humatch to design sequences that pass all CNN thresholds of 0.95 in fewer edits than otherwise possible at the expense of lower alignment with experimental endpoints. Table S5 shows Humatch's performance when humanising 25 precursor therapeutics using GL targets of 0.40 vs 0.38 (approximately corresponding to the lower 10th percentile of GL scores achieved by CSTs, see Figure S6). Fewer edits are made to both heavy and light chains using the lower GL score, but the overlap of Humatch's designs with experimental designs also falls.

| GL target | H overlap | L overlap | H edit | L edit |
|---|---|---|---|---|
| 0.40 | 0.77 | 0.82 | 20.6 | 13.0 |
| 0.38 | 0.76 | 0.79 | 18.4 | 11.3 |

Table S5: Comparison of Humatch's designs when humanising the same 25 precursor therapeutics using different initial germline-likeness targets. 'H edit' and 'L edit' are the mean edit distances of the humanised therapeutics from their corresponding precursor sequences. 'H overlap' and 'L overlap' described the mean overlap in mutations made computationally compared to those made experimentally. If all suggested computational mutations were made experimentally, the overlap would be one; if none matched, the overlap would be zero.

### 12 Humatch humanisation of 25 precursor therapeutics allowing CDR mutations

CDR mutations can be allowed by the user in both stages of Humatch's humanisation pipeline: stage 1 - the initial germline-likeness matching, and stage 2 - the subsequent iterative design and selection of single-point mutants. By default, Humatch does not allow CDR mutations in either stage to increase its speed (especially in stage 2, as fewer single-point variants require screening). Table S6 also shows that disallowing CDR mutations results in designs with greater experimental overlap by examining Humatch's performance when humanising 25 precursor therapeutics with known therapeutic endpoints. Nevertheless, if users do wish to allow mutations within the CDR regions, Figure S9 shows that these are likely to be small in number and located near the CDR anchors.

| Stage 1 | Stage 2 | H overlap | L overlap | H edit | L edit |
|---|---|---|---|---|---|
| Disallow | Disallow | 0.77 | 0.82 | 20.6 | 13.0 |
| Allow | Disallow | 0.73 | 0.80 | 21.2 | 13.0 |
| Disallow | Allow | 0.75 | 0.76 | 20.4 | 13.4 |
| Allow | Allow | 0.72 | 0.75 | 20.9 | 13.2 |

Table S6: Comparison of Humatch's designs when humanising the same 25 precursor therapeutics either allowing or disallowing CDR mutations in the two stages of Humatch's humanisation protocol. 'Stage 1' refers to the initial germline-likeness matching, and 'Stage 2' refers to the iterative design and selection of single-point variants. 'H edit' and 'L edit' are the mean edit distances of the humanised therapeutics from their corresponding precursor sequences. 'H overlap' and 'L overlap' described the mean overlap in mutations made computationally compared to those made experimentally. If all suggested computational mutations were made experimentally, the overlap would be one; if none matched, the overlap would be zero.

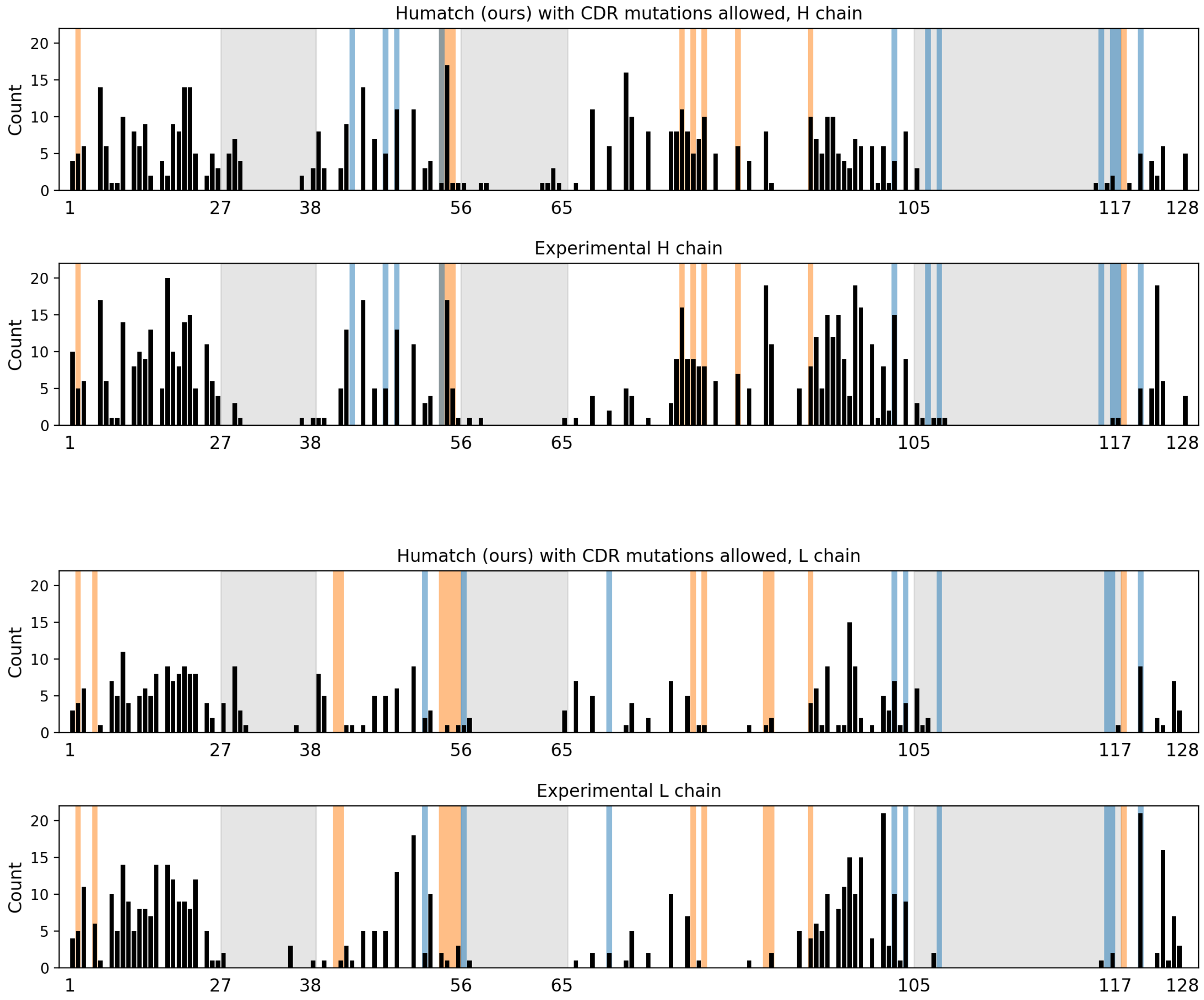


Figure S9: A comparison of Humatch and experimental mutation profiles for 25 precursor therapeutics, separated by heavy and light chains. Here, Humatch was run disallowing CDR mutations (top) and while allowing CDR mutations in both stages of its humanisation protocol (bottom). The black bars show the number of mutations made at each IMGT sequence position across the 25 therapeutics. CDR regions are shaded in grey, Vernier zone residues in orange, and interface residues in blue.

### 13 Nearest training data matches to 25 experimental endpoints

To check if Humatch simply memorised its training data and pushed designs towards these sequences during humanisation, we compared the 25 experimental endpoints to their closest training/validation matches (Figure S10). The closest sequence matches were identified using KASearch with its pre-aligned version of OAS[5] containing 2.4 billion sequences. We returned the closest 20 entries for every experimental heavy and light endpoint and then compared these against our training and validation data. As we did not include all OAS sequences in our training, some therapeutics returned no matches for all 20 hits (grey boxes in Figure S10). When hits were found, they were found to be distant, with mean heavy and light edit distances of 21 and 11 between the experimental endpoints and the closest training sequences. For comparison, we also include the edit distances between our designs and all experimental endpoints, and our designs and the closest experimental training hits in Figure S10. We see that our designs are closer to the experimental designs than their closest training dataset matches, indicating that Humatch does not simply push designs towards sequences it has observed.

| Therapeutic | H-edit (exp-ours) | L-edit (exp-ours) | H-edit (exp-closest) | L-edit (exp-closest) | H-edit (ours-closest) | L-edit (ours-closest) |
|---|---|---|---|---|---|---|
| AntiCD28 | 20 | 10 | 20 | 19 | 29 | 23 |
| Bevacizumab | 12 | 7 | 29 | 8 | 27 | 15 |
| Campath | 16 | 8 | | 10 | | 17 |
| Certolizumab | 13 | 11 | 25 | 13 | 28 | 17 |
| Clazakizumab | 11 | 6 | | 14 | | 19 |
| Crizanlizumab | 15 | 12 | 23 | 11 | 25 | 19 |
| Eculizumab | 16 | 4 | 22 | | 28 | |
| Etaracizumab | 16 | 20 | 15 | 13 | 22 | 21 |
| Herceptin | 12 | 12 | | 10 | | 17 |
| Idarucizumab | 13 | 10 | 24 | | 29 | |
| Ixekizumab | 21 | 10 | 19 | 11 | 26 | 15 |
| Ligelizumab | 16 | 14 | 24 | 11 | 30 | 16 |
| Lorvotuzumab | 6 | 8 | | 8 | | 16 |
| Mogamulizumab | 9 | 9 | 20 | 14 | 25 | 21 |
| Omalizumab | 15 | 11 | 25 | 12 | 29 | 20 |
| Palivizumab | 11 | 11 | | | | |
| Pembrolizumab | 18 | 12 | | | | |
| Pertuzumab | 13 | 10 | 23 | 12 | 24 | 16 |
| Pinatuzumab | 13 | 9 | 22 | | 24 | |
| Refanezumab | 11 | 11 | 16 | 4 | 25 | 15 |
| Reslizumab | 16 | 11 | | 11 | | 16 |
| Rovalpituzumab | 14 | 14 | | 10 | | 18 |
| Solanezumab | 14 | 6 | 12 | 9 | 20 | 15 |
| Talacotuzumab | 23 | 10 | | 8 | | 16 |
| Tocilizumab | 15 | 7 | 20 | 9 | 21 | 16 |

Figure S10: Edit distances between our designs, experimental (exp) designs, and the training/validation sequences closest to the experimental designs for 25 humanised therapeutics. Results are split by heavy (H) and light (L) chains. Smaller edit distances are coloured in green and larger distances in red. The closest sequences were identified using KASearch and its pre-aligned version of OAS. As Humatch was not trained on all of OAS, some closest hits found using were not in the training data - these are shown as grey boxes.

### 14 Examining whether different tools humanise 25 precursor therapeutics between genes

Figure 4 in the main text shows no evidence of any tool humanising ‘between’ genes according to Humatch’s CNN-H and CNN-L classifiers i.e. no two heavy/light V-genes simultaneously show as ‘hot’ in the heatmap. Figure S11 does not provide direct evidence for this either, but it does show that some tools’ designs have germline content less clearly associated with only one V-gene. To avoid bias in this analysis, we did not use Humatch’s classifications and instead only considered germline-likeness (GL) scores that measure how often amino acids are observed at different positions in sequences from OAS. First, we obtained the GL scores for every V-gene for all 25 designs from each tool. The first and second-highest GL scores for each sequence were then selected and their difference was calculated (ΔGL). Larger ΔGL values occur for sequences that share many commonalities with one V-gene as they are less able to match others. Smaller values indicate a sequence is similar to at least two V-genes. We observed that designs from Hu-mAb and AbNatiV generally had lower ΔGL scores. Designs from Sapiens and Humatch had higher ΔGL scores similar to those of experimentally humanised sequences. However, Humatch achieved these higher scores in fewer mutations than Sapiens and did so while targeting the same genes as experiments, unlike Sapiens.

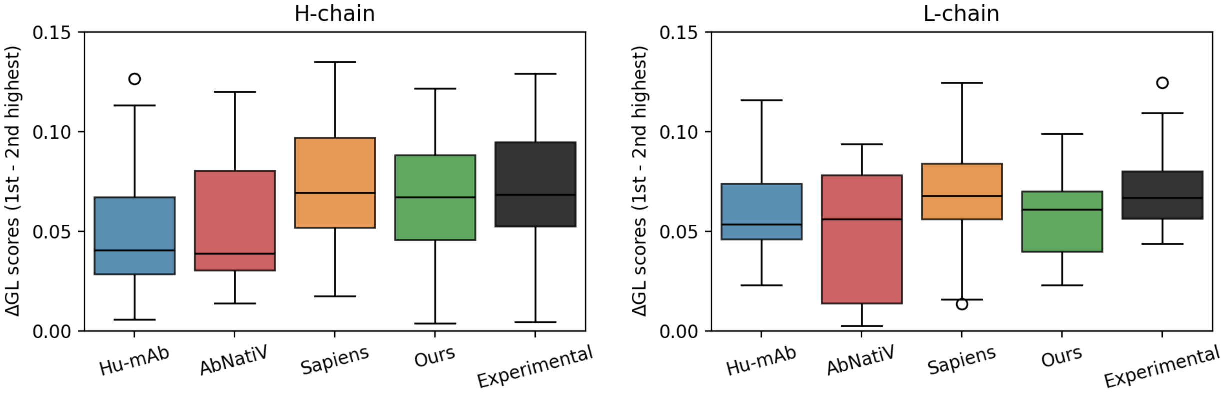


Figure S11: Boxplots showing the differences in the highest and second-highest germline-likeness (GL) scores for 25 humanised heavy and light designs from Hu-mAb, AbNatiV, Sapiens, Humatch, and experimental efforts.

### 15 Humatch classifies more ‘mixed-gene’ sequences as non-human than other tools

To test whether each humanisation tool can successfully identify ‘mixed-gene’ designs as non-human, we created a dataset of artificial mixed heavy V-gene sequences e.g. first half HV1, second half HV3 (see Figure S12). We created this dataset by sampling heavy V-gene sequences from Humatch’s test set. We split sequences in the middle of the CDR2 (IMGT position 61E) as this is approximately the mid-point of the V-gene encoded region (IMGT positions 1 to 106).

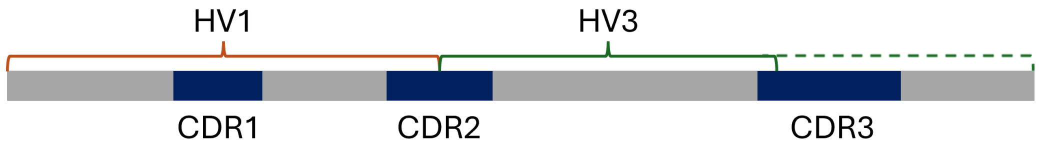


Figure S12: Overview of our mixed-gene designs - the start of the sequence, up to the middle of the CDR2 loop, is taken from one V-gene, while the rest of the sequence is from another.

As sequences were split in the middle of the CDR2, we only re-joined sequence halves that shared original CDR2 lengths. All heavy V-genes are dominated by a single-length CDR2 due to their germline origins: HV1&3&5&7 CDR2s are largely length-eight, HV2&4 are length-seven, and HV6 is length-nine. We ignored sequences with less common CDRH2 lengths resulting from insertions/deletions and all HV6 sequences, instead joining only those from HV1&3&5&7 together, and HV2&4 together. The resulting dataset - both the ∼57k single gene sequences sampled from Humatch’s test set and the ∼122k mixed-gene creations - can be found at doi.org/10.5281/zenodo.13764770

Humatch, using a CNN-H cutoff of 0.95 (corresponding to near-perfect precision and 95.3% recall, see Table S2), successfully classified 51% of the 122k mixed-gene designs as non-human. OASis does not recommend a classifier cut-off in their paper so we instead calculated OASis Identity scores for our 57k single-gene sequences and determined the threshold (0.465) that also gave 95.3% recall. We found that only 1% of mixed-gene designs (fewer than true human sequences) fell below this threshold.

A similar process was followed for AbNatiV, giving a 95.3% recall threshold of 0.736. This cut-off removed 28% of mixed-gene designs, more than OASis but fewer than Humatch. AbNatiV do however recommend a cut-off of 0.8 in their paper. This cut-off gives a lower recall of 83.3% on our 57k single-gene sequences but removes 67% of mixed-gene designs. At 83.3% recall (CNN-H cut-off of 0.99994), Humatch successfully classifies 94% of mixed-gene designs as non-human.

Finally, Hu-mAb was used to score the single and mixed gene sequences. Hu-mAb includes its own classification thresholds for each gene and, coincidentally, these provided an overall recall of 95.3% on our single-gene designs too. Using these default thresholds, Hu-mAb scored only 8% of mixed-gene designs as non-human. Furthermore, we found that 59% of mixed-gene designs were ranked as human according to multiple V-gene classifiers. This in itself is not enough to determine if a sequence is mixed-gene though as Hu-mAb also ranked 24% of the 57k single-gene designs as human according to multiple V-gene classifiers.

Note - OASis and Humatch scored all mixed-gene sequences as OASis does not number sequences and the sequences were pre-aligned for use by Humatch. However, Hu-mAb and AbNatiV both used ANARCI[6] to number the sequences before scoring. ANARCI failed to number ∼2.2% of mixed-gene sequences so these were not scored by either tool.

### 16 Humanising 1,000 non-human sequences

In the main text, we report the mean heavy and light edit distances and run times for the humanisation of 1,000 non-human sequences. Figure S13 shows the full distribution of values from this process. The majority of sequences are humanised in less than a minute and fewer than 40 edits.

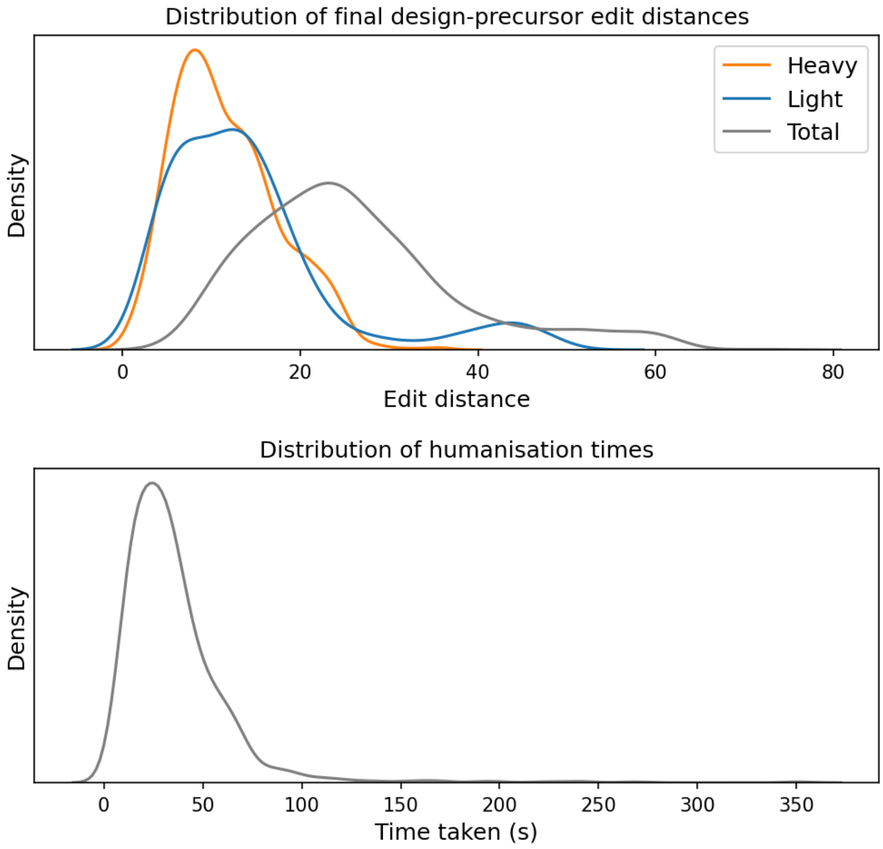


Figure S13: Distribution of heavy and light edit distances of 1,000 Humatch designs from their precursor non-human sequences, and the times taken to reach these designs.

### 17 Hu-mAb, AbNatiV, and Sapiens settings

We ran Hu-mab, AbNatiV, and Sapiens with default and/or paper-recommended parameters. Hu-mAb was run from SAbBox, a vagrant VirtualBox containing many SAbPred[7] tools. Hu-mAb's 25 experimental heavy/light target scores and 211 ADA-therapeutic scores were obtained using e.g.

```
singularity exec /path/to/sabbox.sif Hu-mAb --h_seq <H_seq> --name
<therapeutic> --score_only
```

Sequences were then humanised using e.g.

```
singularity exec /path/to/sabbox.sif Hu-mAb --name <therapeutic>
--h_seq <H_seq> --l_seq <L_seq> --vgene_h <H_gene> --vgene_l
<L_gene> --threshold_h <H_target> --threshold_l <L_target> -v
```

AbNatiV's humanisation code was cloned from gitlab.developers.cam.ac.uk/ch/sormanni/abnativ. Sequences were scored using (for `VH`, `VKappa`, and `VLambda`) e.g.

```
abnativ score -nat VLambda -i <VL_fasta> -odir <VL_odir> -oid test
-align
```

Sequences were humanised using e.g.

```
abnativ hum_vh_vl -i_vh <H_seq> -i_vl <L_seq> -odir </out/dir> -oid
test_vh_vl
```

Sapiens and OASis were run from the web interface - biophi.dichlab.org. Sequences were input in fasta format, IMGT definitions were used for numbering and CDR definitions, and a relaxed OASis prevalence threshold of 10% was used for both scoring and humanisation. Sapiens was used for humanisation and three humanisation iterations were permitted. The CDRs were not allowed to be humanised.
